# Multimodal Virtual Screening Identifies Candidate Inhibitors of Polyploid Giant Cancer Cells Across Breast Cancer Subtypes

**DOI:** 10.64898/2026.11.22.767421

**Authors:** Huikang Ye, Hsi-Chun Wang, Guanyang Wang, Pei-Ru Lin, Ying-Ju Lai, Chien-Hung Shih, Yushu Ma, Hsiao-Chun Chen, Yuan Zhang, Jinxiong Cheng, Omar Baltagi, Yu-Chiao Chiu, Yu-Chih Chen

## Abstract

Polyploid giant cancer cells (PGCCs) are associated with treatment resistance and tumor recurrence yet identifying compounds that selectively suppress these cells remains challenging. Here, we developed a multimodal machine-learning framework to prioritize PGCC-suppressing compounds by integrating chemical fingerprints, drug-induced cellular morphology, transcriptomic perturbation profiles, and biologically informed molecular annotations. Using screening data from four breast cancer cell lines, we systematically evaluated feature selection, drug representations, and ensemble learning strategies. PGCC-guided feature selection reduced morphology dimensionality by 93.7% while identifying cell-cycle-enriched gene sets, linking predictive features to biologically relevant processes. An ensemble integrating complementary predictors achieved a mean Pearson correlation of 0.403 and an enrichment factor of 5.46 among the top 5% of ranked compounds. In held-out screening data, the ensemble recovered 27.9% of experimentally identified hits, representing approximately 5.6-fold and 2.5-fold improvements over random selection and the best individual predictor, respectively. We subsequently prioritized 21,367 compounds from an external library for experimental evaluation. Experimental testing identified six candidates that preferentially suppressed PGCCs in at least three paclitaxel-treated cell lines. Notably, AZD-8055 and voxtalisib reduced PGCC abundance by at least 70% across all four cell lines. These results establish a multimodal computational-experimental strategy for phenotype-directed drug discovery and provide new candidates for targeting treatment-induced PGCCs.

## Introduction

Breast cancer remains a major cause of cancer-related death among women worldwide,^1^ and the persistence of treatment-resistant cancer cells limits durable disease control.^2^ Genomic analyses have identified whole-genome doubling in approximately half of breast tumors in the cohorts examined,^3,4^ and its presence has been associated with poorer survival in advanced cancers.^5^ In a small breast-cancer biopsy study, near-triploid DNA content was associated with weak or absent response to neoadjuvant chemotherapy.^6^ At the cellular level, PGCCs are enlarged cells with increased genomic content, and their abundance in breast tumors has been associated with lymph-node metastasis.^7^ PGCCs can survive treatment-associated stress^8–11^ and generate proliferative progeny with stem-like properties, providing a potential route to tumor repopulation.^12–14^ Paclitaxel is widely used in breast-cancer chemotherapy,^15^ and experimental studies have shown that it can induce the formation of PGCCs whose progeny are more resistant to paclitaxel.^16^ Together, these findings underscore the need to identify therapeutics that selectively eliminate PGCCs, both as a standalone strategy and in combination with chemotherapy, to limit the survival and subsequent regrowth of treatment-resistant cancer cells.

Identifying compounds that suppress PGCCs requires their selective quantification within heterogeneous cell populations, as reductions in overall cell viability may mask the persistence of a smaller, treatment-resistant PGCC population.^17^ Flow cytometry is the gold standard for identifying PGCCs based on DNA content and cell size; however, its requirement for cell harvesting and resuspension from individual wells limits its scalability for high-throughput screening. We previously established a high-throughput imaging-based workflow that separately quantifies viable PGCCs and non-PGCCs in each well.^18^ Even with this workflow, testing every candidate, dose and combination would require an impractical number of experiments. In silico prediction offers a way to prioritize these experiments by ranking compounds before testing. Large pharmacogenomic studies have linked molecular characteristics of cancer cell lines to drug sensitivity,^19^ enabling models that combine genomic, transcriptomic, chemical and cell-line features to estimate response.^20–23^ For PGCC screening, however, these models must predict suppression of the PGCC population, rather than overall cell viability. Cross-study analyses show that predictive performance can change between screening datasets,^24,25^ reinforcing the need to evaluate PGCC prediction using population-specific response measurements.

While our previous study established the feasibility of PGCC prediction, it lacked PGCC-informed feature selection and evaluated an ensemble limited to chemical fingerprints and drug-description embeddings.^26^ That ensemble used linear regression to combine predictions from a fingerprint-based gradient-boosting model and a drug-description-based support vector machine. Trained exclusively in MDA-MB-231, it did not model differences in drug sensitivity across cell lines. Extending this approach requires broader drug profiles and prediction across multiple cell lines. Transcriptomic profiles measure gene-expression changes, whereas Cell Painting captures alterations in cellular and organelle morphology.^27–29^ Feature selection can prioritize PGCC-relevant measurements from these high-dimensional profiles, while curated target/pathway information^30^ and language-model-derived annotations^31,32^ provide additional pharmacological context. Prior studies have demonstrated that imaging profiles can predict activity in other assays^33^ and that integrating chemical, morphological, and transcriptional features broadens prediction across assays.^34^ Cell-line molecular characteristics describe the biological background in which drug responses are measured.^35^ The challenge is to select PGCC-relevant features and combine complementary predictors to prioritize active compounds across breast cancer cell lines.

Building on our previous screening and prediction studies,^18,26^ we sought to overcome these limitations by developing a multimodal framework to identify PGCC-suppressing compounds across four breast cancer cell lines of different molecular subtypes. By integrating complementary drug profiles, the framework aims to improve compound prioritization across cell lines. At the same testing budget, retrospective evaluation recovered 5.6 times as many hits as random selection and 2.5 times as many as the best individual predictor. We further applied the framework to prioritize previously untested compounds, identifying six candidates that strongly suppressed PGCCs in paclitaxel-treated cultures while preferentially sparing non-PGCCs. This approach provides a strategy for leveraging complementary biological and chemical information to accelerate the discovery of therapeutics targeting PGCCs.

## Results

### Multimodal framework for PGCC prediction

To develop models that predict PGCC drug responses and prioritize candidate compounds for experimental validation, we used our published screen of 2,691 clinical-stage and FDA-approved compounds. 18 The four breast cancer cell lines represented major molecular subtypes: triple-negative SUM159 and MDA-MB-231, hormone-receptor-positive MCF7, and HER2-amplified SK-BR-3.36 The framework used drug profiles together with baseline molecular features for each cell line (Fig. 1a, b). Drug profiles comprised six information families: (1) chemical fingerprints; (2) curated target and pathway annotations; (3) large language model (LLM)-derived annotations and biological ratings; (4) transcriptomic perturbation profiles; (5) Cell Painting morphology profiles; and (6) language-model embeddings of chemical structures or drug descriptions. Each family contains one or more feature sets representing distinct aspects of the compounds. Cellular context included baseline mutation, copy-number, gene-dependency, and gene-expression features for each cell line. Supplementary Data 1 summarizes the feature sets and their sources.

**Figure 1.**
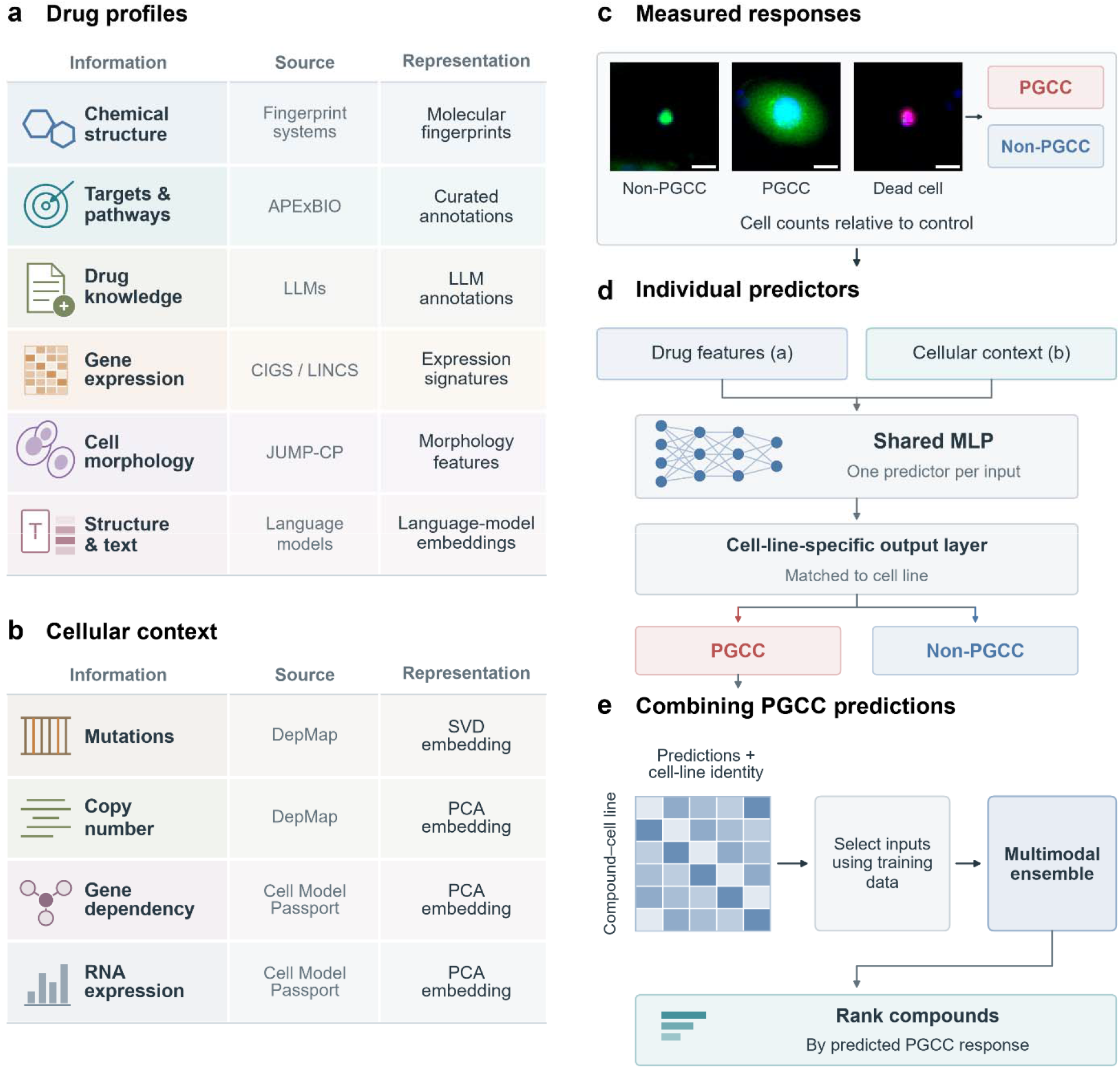
Prediction framework. Multimodal framework for PGCC drug prediction. a, Six drug-information families describe chemical structure, curated targets and pathways, LLM-derived annotations, transcriptomic perturbations, cell morphology and language-model embeddings of structures or drug descriptions. Sources and input types are listed for each family. b, Mutation, copy-number, gene-dependency, and RNA-expression profiles provide cellular context. Mutation features are reduced by singular value decomposition (SVD), and the other context features by principal component analysis (PCA). c, SUM159 single-cell images show a non-PGCC, a PGCC and a dead cell (paclitaxel). Blue, Hoechst-stained nuclei; green, live-cell staining; red, dead-cell staining. Scale bars, 20 µm. The two outputs are the viable PGCC and non-PGCC counts, each expressed as a log2 ratio relative to controls. d, For each input feature set, raw or learned drug representations are combined with cellular context in an individual MLP. Each input passes through shared hidden layers and then through the output layer matched to its cell line to predict PGCC and non-PGCC responses. e, Individual PGCC predictions and cell-line identity form the inputs to the multimodal ensemble. Predictor selection uses training data, and TabPFN combines selected predictions to rank compounds within each cell line. Rows represent compound–cell-line pairs.

The screen quantified viable PGCC and non-PGCC populations separately, providing two response outcomes illustrated by the single-cell examples in **Fig. 1c**. Counts were normalized to controls and expressed on a log2 scale, with lower values indicating greater suppression of the corresponding population. We trained individual predictors using each drug-feature input together with the same cellular-context features. Each multilayer perceptron (MLP) used shared hidden layers to learn common drug-response patterns across cell lines, followed by cell-line-specific output layers to accommodate differences in their responses (**Fig. 1d**). Individual predictors were evaluated by repeated compound-level cross-validation. We then integrated their PGCC predictions into a multimodal ensemble to capture complementary information across feature families (**Fig. 1e**). Ensembles were evaluated using ten matched splits, with approximately 80% of compounds assigned to training and 20% to testing. All measurements for a compound were kept together in each ensemble split. The ensemble produced a separate list of compounds to prioritize for each cell line. Adding cellular context provided variable, input-dependent benefits, improving predictive performance for certain information families while offering limited or no gains for others. (**Supplementary Fig. 1**).

To determine whether learned representations improved downstream prediction, we compared alternative numerical representations of drug profiles. **Figure 2** illustrates these approaches using molecular fingerprints. Raw fingerprints encode molecular substructures as binary features (**Fig. 2a**). MLP autoencoders (MLP-AEs) learn compact representations by reconstructing the input fingerprints (**Fig. 2b**), ^37^ whereas variational autoencoders (VAEs) additionally regularize the latent representations toward a specified distribution (**Fig. 2c**). ^38^ Contrastive learning trains an encoder to represent perturbed versions of the same compound similarly (**Fig. 2d**).^39^ We also applied these approaches to other drug inputs. For each input, we used the same type of MLP response predictor to test whether encoding the features improved prediction over using them directly (**Fig. 2e**).

**Figure 2.**
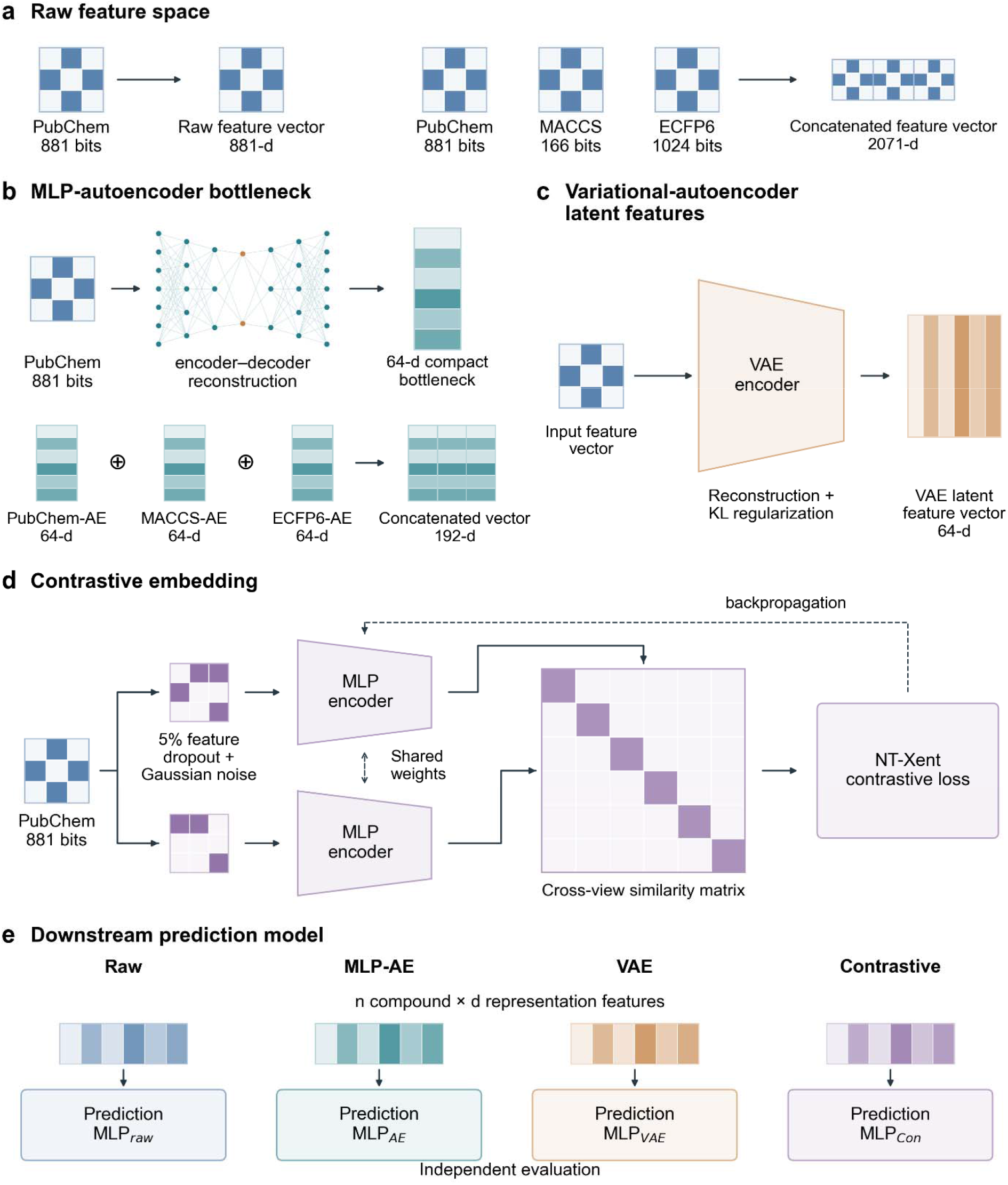
Fingerprint representations. Raw and learned fingerprint representations for PGCC response prediction. a, PubChem, MACCS and ECFP6 fingerprints contain 881, 166 and 1,024 bits, respectively. b, Separate autoencoders learn 64-dimensional representations by reconstructing each fingerprint. Concatenating the three 64-dimensional vectors yields a 192-dimensional representation. c, The fingerprint VAE uses a 64-dimensional latent representation with reconstruction and Kullback– Leibler regularization. d, Feature dropout and Gaussian noise generate two views of the same input. The contrastive objective encourages similar representations of these paired views relative to other compounds. e, Raw and learned representations are evaluated using individual MLP predictors. Fingerprints illustrate these methods; contrastive encoders were also applied to other drug-feature inputs.

We evaluated predictive performance using two complementary objectives: response estimation and compound prioritization. Pearson correlation measured agreement between predicted and observed response values, whereas enrichment factor at a 5% testing budget (EF@5%) assessed the recovery of strongly suppressing compounds. A hit was defined as a compound reducing measured PGCC counts by at least 70% (count ratio ≤0.30). EF@5% compared the hit fraction among the highest-ranked 5% of compounds with that in the full test set. Thus, these metrics assessed both quantitative prediction accuracy and the ability to prioritize active compounds under a limited experimental screening budget.

### Biologically guided features support response prediction

Given the established role of cell-cycle dysregulation in PGCC formation, ^40^ we selected five Aurora kinase inhibitors, three microtubule-targeting agents, and two PARP inhibitors to guide Cell Painting feature selection. Our previous breast cancer study showed that the Aurora kinase inhibitor tozasertib increases PGCC prevalence, ^41^ consistent with reports that Aurora kinase inhibition induces PGCC formation through endoreplication in lung cancer cells. ^42^ Prior studies have also linked microtubule disruption to tetraploidy^43^ and PARP inhibition to PGCC formation.^44^ Because these drug classes are established PGCC inducers, we used the morphological features they elicited as candidate features associated with the PGCC phenotype. To test whether morphological responses shared across the reference treatments could inform PGCC drug-response prediction, we ranked Cell Painting features by the absolute value of their mean change relative to DMSO across the ten compounds (**Fig. 3a**). The largest shifts involved nuclear shape, nuclear-edge DNA intensity, and spatial patterns of RNA and endoplasmic-reticulum staining (**Fig. 3b**). We evaluated models using the top 50, 100, 200, 300, or 500 ranked features under matched training conditions (**Fig. 3c**). Across the four cell lines, the 200-feature model achieved a mean Pearson correlation of 0.191 and an EF@5% of 2.93. Neither metric differed significantly across the feature-set sizes tested, indicating comparable predictive performance over this range. We therefore selected 200 features as a practical model-design choice that maintained predictive performance while limiting input dimensionality, computational burden, and the risk of overfitting.

**Figure 3.**
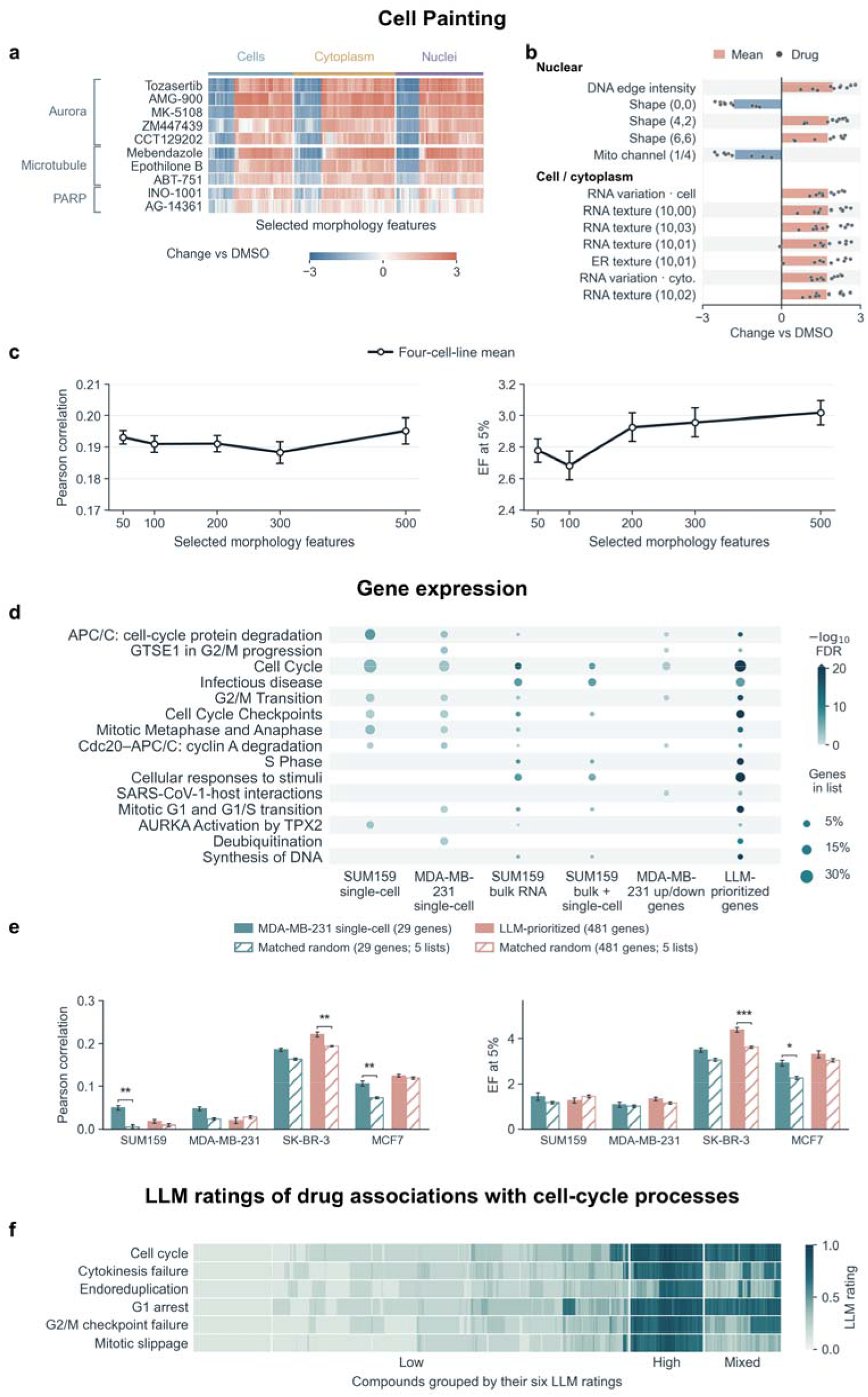
Feature construction and evaluation. Biologically guided features for predicting PGCC response. a, Cell Painting profiles of ten reference compounds. Each value is the replicate-averaged drug profile minus the feature-specific mean from 384 DMSO wells, on a common linear scale from −3 to +3. Rows are grouped by drug mechanisms and columns by compartment. b, The twelve nontechnical measurements with the largest absolute mean shifts, grouped by object mask. Bars show signed means and dots show individual drugs. c, Pearson correlation and EF@5% from pooled out-of-fold predictions using the most altered Cell Painting features. We compared the top 50, 100, 200, 300, and 500 features ranked by the magnitude of their mean change across the ten reference-drug treatments relative to DMSO. The evaluation included 1,586 compounds per cell line. Points and whiskers show mean ± SEM across ten repetitions, each averaging the four cell lines equally. d, Reactome pathway enrichment for the six gene lists, using 3,400 mapped CIGS genes as the background. Dots show Benjamini–Hochberg FDR <0.05 across 6,684 tests. Color shows −log10(FDR), capped at 20. Area gives the percentage of each list assigned to a pathway. e, Performance of contrastive + MLP predictors using the MDA-MB-231 single-cell list (29 genes) and the LLM-prioritized list (481 genes) across four cell lines. Solid bars show each selected list, and adjacent hatched bars show the mean of its five size-matched random lists. Bars and error bars show mean ± SEM across ten repetitions; random-list metrics were averaged within each repetition. Asterisks indicate two-sided paired compound-bootstrap comparisons with the random-list mean (*P <0.05, **P <0.01, ***P <0.001). Exact P-values and paired confidence intervals are in **Supplementary Data 1**. f, GPT-4o ratings of drug associations with six cell-cycle processes. Ward clustering used Euclidean distances across the six ratings. White separators mark the three clusters. The high-rating group had the largest mean across its compounds and all six ratings.

We asked whether genes selected from measured differences between PGCCs and non-PGCCs, or from prior knowledge of PGCC biology, could identify drug-induced expression changes predictive of PGCC response. For transcriptomic perturbation profiles, we evaluated five gene sets derived from single-cell and bulk RNA analyses of PGCCs and non-PGCCs in SUM159 and MDA-MB-231^41^ and one LLM-prioritized set; their definitions are listed in **Supplementary Data 1**. Reactome enrichment identified cell-cycle pathways across the lists (**Fig. 3d**).^45^ To assess predictive information beyond gene-set size, we compared the MDA-MB-231 single-cell list (29 genes) and the LLM-prioritized list (481 genes) with five size-matched random lists each, using matched contrastive encoders and MLP predictors. Both lists gave higher mean Pearson correlation and hit enrichment than their random controls in SK-BR-3 and MCF7 (**Fig. 3e**). In SK-BR-3, the LLM-prioritized list increased Pearson r from 0.194 to 0.220 (P = 0.00162) and EF@5% from 3.64 to 4.38 (P = 0.00050). In MCF7, the MDA-MB-231 single-cell list increased Pearson r from 0.072 to 0.106 (P = 0.00428) and EF@5% from 2.25 to 2.92 (P = 0.0434). These comparisons show that PGCC-guided gene selection can retain predictive information in compact expression profiles, with gains that vary across cell lines.

Alongside the measured expression profiles, we asked whether ratings based on drug mechanisms would group compounds with known effects on cell division. GPT-4o rated each screened compound for its association with six cell-cycle processes, including cytokinesis failure, endoreduplication, and mitotic slippage (**Fig. 3f**). Ward clustering of the six ratings produced three groups. We designated the 340 compounds in the cluster with the highest mean across all six ratings as the high-rating group (mean 0.725 on a 0–1 scale). This group included all 16 compounds with a curated microtubule/tubulin annotation and 18 of the 19 with an Aurora kinase annotation. The clustering of both drug classes in this group is consistent with evidence that perturbing microtubule dynamics or Aurora kinase activity can promote polyploidization.^41–43^

### Comparing prediction algorithms and drug representations

We first compared MLP with random forest (RF), histogram gradient boosting (HGB), and support vector machine (SVM) regression across six input families. MLP had the highest overall held-out Pearson r in each of the six input families (**Fig. 4a–f**). We then evaluated whether reconstruction-or contrastive-learning-based representations improved MLP prediction of PGCC response on held-out compounds compared with the corresponding raw features (**Fig. 4g**). All three encoded representations showed numerically higher mean Pearson correlations, with gains of 0.021–0.043. The improvements were statistically significant for MLP-AE encoding of ECFP6 fingerprints (0.217 versus 0.174, P = 0.0426) and contrastive encoding of curated targets and pathways (0.234 versus 0.214, P = 0.0286). VAE encoding of Cell Painting features also increased mean correlation (0.176 versus 0.150), although this difference did not reach statistical significance (P = 0.0732). Thus, learned representations improved prediction for two of the three inputs tested, while showing a consistent numerical trend across all three.

**Figure 4.**
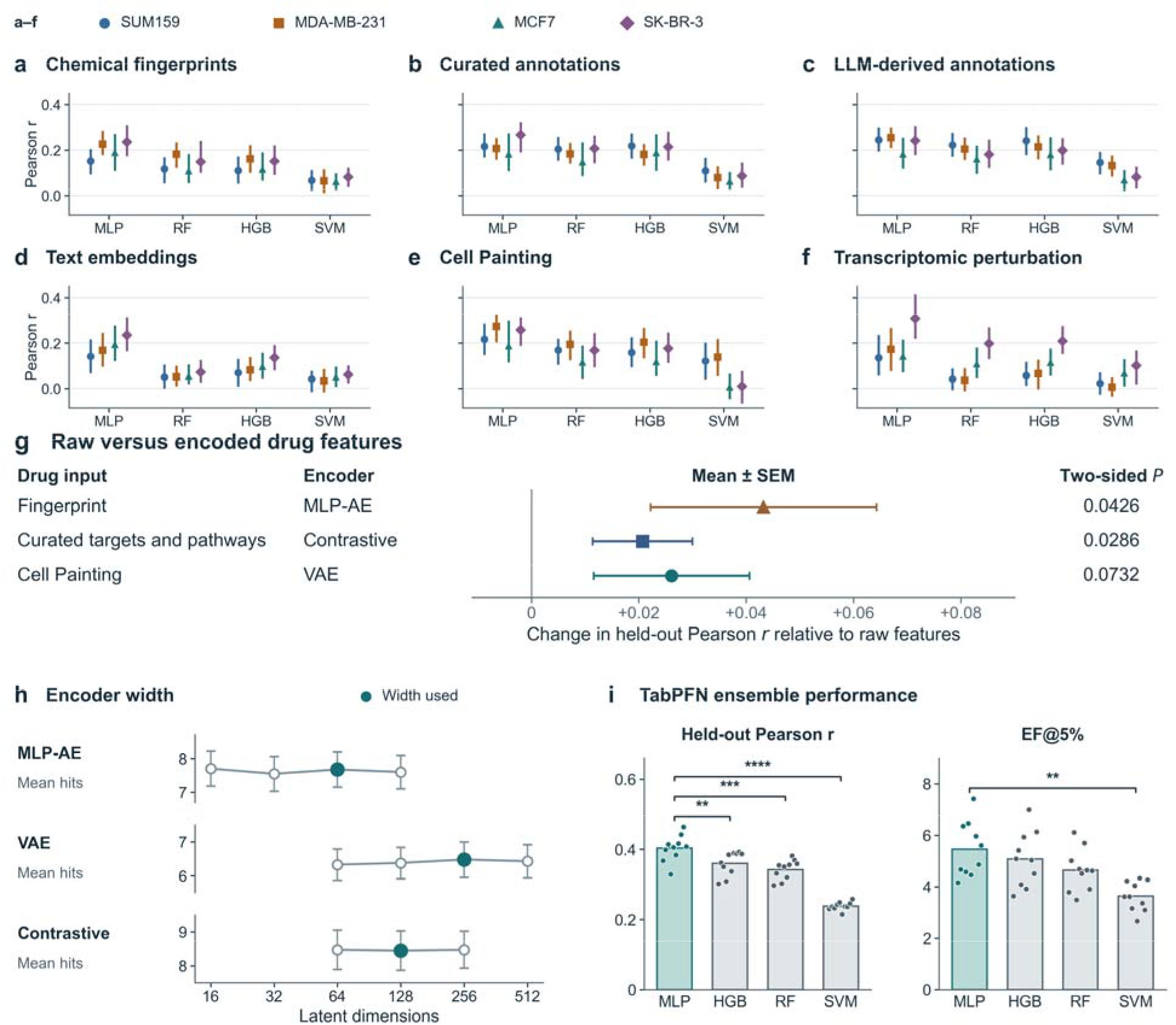
Representation and model comparisons. Drug representations and prediction algorithms for PGCC response. a–f, Base-algorithm performance for chemical fingerprints (a), curated annotations (b), LLM-derived annotations (c), text embeddings (d), Cell Painting (e) and transcriptomic perturbations (f). Points and vertical lines show medians and interquartile ranges of 400 fold scores per cell line and algorithm, pooled from the two highest-ranked feature sets per family and their representations (Methods). g, Three selected examples of PGCC response prediction using encoded drug features. Points show the change in held-out Pearson correlation relative to an MLP predictor using the corresponding raw features. Changes are averaged across four cell lines; horizontal lines show SEM. Fingerprint denotes ECFP6. For ECFP6 and curated annotations, SEMs and two-sided paired t-tests accounted for fold overlap (100 paired folds). For Cell Painting, SEMs and two-sided P-values used a paired compound bootstrap across ten matched holdouts. h, Mean number of hits per 27-compound test list at each encoder width. Points and error bars show mean ± SEM across ten splits, each averaging four cell lines. Fingerprint MLP-AE widths varied together across ECFP6, MACCS and PubChem inputs and their selected combinations. The VAE and contrastive experiments varied the SUM159 bulk gene-expression and Cell Painting predictors, respectively. Filled circles mark the widths used for screening. Hits had a measured PGCC count ratio ≤0.30. i, Held-out Pearson correlation and EF@5% for TabPFN ensembles using MLP, HGB, RF or SVM base predictors. Each ensemble selected inputs from the same 82 candidates using training data. Points show four-cell-line means for individual compound splits, and bars show means across ten matched splits. Brackets show pairwise comparisons of MLP with each alternative using two-sided, overlap-corrected paired t-tests (*P < 0.05, **P < 0.01, ***P < 0.001, ****P < 0.0001).

We evaluated different latent dimensions to identify practical settings that balanced screening performance and model complexity (**Fig. 4h**). Each comparison used a top 5% testing budget across four cell lines and ten matched compound splits, yielding 40 lists of 27 compounds per width. For the combined ECFP6, MACCS, and PubChem MLP-AE representations, the retained width of 64 dimensions recovered a mean of 7.68 hits per list, comparable to the maximum of 7.70 at 16 dimensions. For the SUM159 bulk gene-expression VAE, 256 dimensions yielded the highest mean recovery (6.48 hits per list) among the four widths tested. For contrastive Cell Painting, 128 dimensions recovered 8.45 hits per list, nearly matching the maximum of 8.48 at both 64 and 256 dimensions. Thus, the selected widths maintained hit recovery close to the best observed performance while limiting input dimensionality and computational burden. These settings were retained as engineering choices for subsequent screening, rather than as statistically optimal dimensions.

Different drug families capture distinct feature representations and may contribute unequally to predicting PGCC responses. Therefore, integrating these complementary representations with appropriate feature selection and weighting is important for improving predictive performance. To achieve this, we trained ensemble models to integrate predictions from individual drug families with cell-line identity. We compared a pretrained tabular model (TabPFN),^46,47^ three boosted-tree methods and two-layer stacking^48^ with simple averaging and the best individual predictor (**Tables 1–2**). All four base learners shared the same pool of 82 candidate inputs and the same ten compound splits. TabPFN gave the highest mean held-out Pearson r for each base learner. With MLP base predictors, TabPFN improved both Pearson correlation (0.403 versus 0.359, P = 0.0179) and EF@5% (5.46 versus 4.55, P = 0.0128) over averaging the same predictions.

**Table 1.** Pearson correlation of combination methods and comparators for four separately evaluated base algorithms. Each column evaluates one base algorithm using 82 shared candidate input identifiers and ten repeated compound-level train–test splits. Preprocessing followed the recorded settings for each algorithm (**Supplementary Computational Methods**). Rows compare five learned methods with the best individual predictor and simple averaging. Test compounds were held out at the ensemble stage (Methods). Each split summary gives equal weight to the four cell-line correlations; values are the mean ± sample SD of those summaries across splits. Simple averaging uses each algorithm’s TabPFN-selected log2 predictions with training-median imputation and no cell-line indicators. Bold marks the highest mean within each row. The best individual predictor was selected using outer-training outcomes. For TabPFN versus averaging, P values were 0.0179, 4.49 × 10, 4.06 × 10 and 4.07 × 10 for MLP, HGB, RF and SVM, respectively.

| **Method** | **MLP** | **HGB** | **RF** | **SVM** |
|---|---|---|---|---|
| Best individual predictor | **0.263 ± 0.056** | 0.176 ± 0.043 | 0.138 ± 0.022 | 0.131 ± 0.017 |
| Simple average | **0.359 ± 0.024** | 0.312 ± 0.037 | 0.295 ± 0.028 | 0.048 ± 0.035 |
| CatBoost | **0.380 ± 0.030** | 0.324 ± 0.037 | 0.312 ± 0.033 | 0.210 ± 0.021 |
| XGBoost | **0.376 ± 0.031** | 0.310 ± 0.045 | 0.308 ± 0.040 | 0.202 ± 0.019 |
| LightGBM | **0.354 ± 0.032** | 0.309 ± 0.041 | 0.302 ± 0.041 | 0.206 ± 0.024 |
| TabPFN | **0.403 ± 0.038** | 0.360 ± 0.034 | 0.342 ± 0.028 | 0.238 ± 0.012 |
| Two-layer stacking | **0.382 ± 0.033** | 0.324 ± 0.031 | 0.322 ± 0.038 | 0.222 ± 0.021 |

**Table 2.** Enrichment for strong PGCC suppression using combination methods and comparators evaluated separately for four base algorithms. EF@5% is the hit fraction among the top 5% of predictions divided by the hit fraction in the corresponding cell-line test set; a hit has a measured PGCC count ratio ≤0.30. Each test set contributed 27 selected compounds. Values are mean ± sample SD across ten splits after equally averaging the four cell-line EF values. Candidate inputs, averaging, bold formatting and evaluation design are as in Table 1. For TabPFN versus averaging, P values were 0.0128, 0.271, 0.208, and 0.00702 for MLP, HGB, RF, and SVM, respectively.

| **Method** | **MLP** | **HGB** | **RF** | **SVM** |
|---|---|---|---|---|
| Best individual predictor | **3.27 ± 0.72** | 2.89 ± 0.78 | 2.41 ± 0.60 | 2.82 ± 0.58 |
| Simple average | 4.55 ± 0.92 | **4.81 ± 1.11** | 4.07 ± 0.92 | 1.60 ± 0.77 |
| CatBoost | **5.06 ± 1.10** | 4.80 ± 1.07 | 4.18 ± 1.21 | 3.71 ± 1.03 |
| XGBoost | **5.09 ± 0.87** | 4.60 ± 1.14 | 4.18 ± 0.86 | 3.22 ± 0.97 |
| LightGBM | **4.74 ± 1.08** | 4.57 ± 1.11 | 4.08 ± 0.96 | 3.27 ± 0.94 |
| TabPFN | **5.46 ± 1.07** | 5.07 ± 1.07 | 4.65 ± 0.82 | 3.64 ± 0.57 |
| Two-layer stacking | **5.19 ± 0.91** | 4.66 ± 1.12 | 4.23 ± 1.15 | 3.34 ± 0.85 |

With TabPFN as the fixed modality blender, the MLP-based ensemble had a higher held-out Pearson correlation than the HGB, RF and SVM ensembles in all ten splits (all P < 0.006, **Fig. 4i**). At the 5% screening budget, the MLP-based ensemble achieved higher hit enrichment than the SVM ensemble (5.46-fold versus 3.64-fold, P = 0.0078). Enrichment was intermediate for HGB (5.07-fold) and RF (4.65-fold). We therefore used TabPFN to combine MLP predictions for screening.

### The multimodal ensemble improves hit recovery and cell-line ranking

At a fixed screening budget, the multimodal ensemble recovered more measured hits than the best individual predictor selected on training data. The best individual predictor was selected from 91 individual MLP predictors spanning six information families. These candidates used different input features and the evaluated raw-feature, MLP-AE, VAE, and contrastive representations. For each split, we selected the predictor with the highest Pearson correlation between its out-of-fold predictions and measured responses in the training partition, pooling the four cell lines. We then evaluated the selected predictor and the multimodal ensemble on the same held-out compounds at the same testing budget. Averaged across four cell lines and ten matched test splits, the ensemble recovered 27.9% of measured hits at a 5% testing budget, compared with 11.3% for the best individual predictor and 5.0% expected for random selection (**Fig. 5a**). The gain over the individual predictor was 16.6 percentage points (95% CI 12.4–21.4, *P* = 0.000020), or 2.5-fold greater recovery. At 1% and 10% budgets, the ensemble recovered 9.7% and 38.3% of hits, compared with 4.2% and 16.7% for the individual predictor. A separate refit that excluded test compounds throughout base-model training retained fourfold hit enrichment (**Supplementary Table 1**).^49^ Across ranking thresholds, the ensemble also achieved higher average precision (0.276 versus 0.111, **Fig. 5b**). Each model ranked the drugs by how strongly it predicted they would suppress PGCCs (**Fig. 5c**). For drugs in the ensemble’s top 10%, measured PGCC counts were 69% of control, compared with 102% of control for drugs in the individual predictor’s top 10% (*P* = 0.000020). For drugs in the bottom 10%, measured counts were 220% of control for the ensemble and 150% for the individual predictor. The ensemble thus placed drugs with stronger measured suppression near the top of its ranking and those with weaker effects near the bottom.

**Figure 5.**
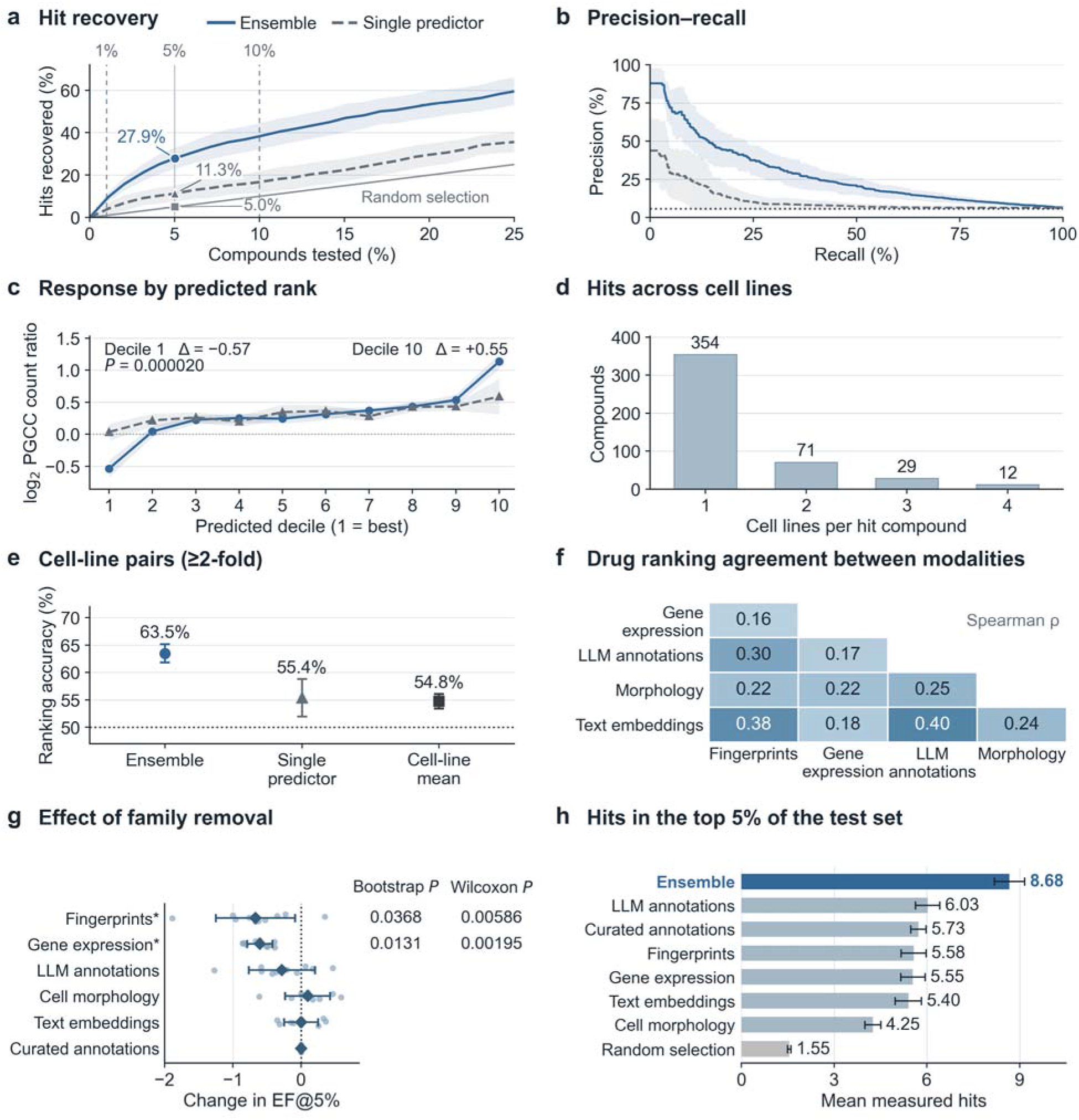
Multimodal prediction of PGCC drug response. Compound prioritization by the multimodal ensemble. A hit had a measured PGCC count ratio ≤0.30. a–c, Blue and dashed gray lines show the ensemble and the individual predictor selected on training data. Bands show SD across ten splits. a, Fraction of measured hits recovered at each screening budget. Vertical lines mark the 1%, 5%, and 10% testing budgets. Points and labels show mean hit recovery at the 5% budget for the ensemble, individual predictor, and random selection. The solid diagonal line shows expected random recovery. b, Precision–recall curves. The dotted line marks hit prevalence. **c, Measured PGCC response across predicted drug ranks.** Each method ranked compounds from the strongest predicted PGCC suppression (decile 1) to the weakest (decile 10). Values below zero indicate fewer PGCCs than in control cultures, and values above zero indicate more. Points show the median measured log2 PGCC count ratio in each decile, averaged across four cell lines and ten test splits. Δ denotes the ensemble minus individual predictor difference. The *P* value shown for decile 1 is from a two-sided paired compound bootstrap. d, Number of cell lines meeting the hit criterion for each active compound. Of 2,690 profiled compounds, 466 were active in at least one line. e, Accuracy in identifying the more sensitive cell line when a drug’s measured responses differed by at least twofold. The dotted line marks 50% accuracy. Whiskers show SD across ten splits. f, Mean pairwise Spearman correlations between the drug rankings predicted by training-selected representatives of the five retained families. Morphology denotes Cell Painting. g, Change in EF@5% after family removal and predictor reselection, relative to the complete ensemble. Points show the ten splits; diamonds and whiskers show mean ± SD. Asterisks mark two-sided compound-bootstrap P < 0.05. For the two marked removals, columns show the bootstrap and additional paired Wilcoxon signed-rank P values. **Supplementary Data 1** lists all test results. Curated annotation removal gave identical predictions. h, Mean measured hits among the top-ranked 5% of each test set (27 compounds), comparing the ensemble with one training-selected predictor per family. Random selection shows the expected hit count at the same budget. Bars and whiskers show mean ± SEM across ten splits, each averaging four cell lines.

Most hits were specific to one cell line. Of the 466 compounds that met the hit threshold in at least one line, 354 (76.0%) met it in one line and 12 (2.6%) met it in all four (**Fig. 5d**). We therefore tested whether the models could identify which of two cell lines was more sensitive to the same drug. When measured responses differed by at least twofold, the ensemble correctly identified the more sensitive line in 63.5% of comparisons, versus 55.4% for the best individual predictor and 54.8% for the cell-line mean baseline (*P* = 0.000020 for each comparison, **Fig. 5e**). These findings suggest potential value in cell-line-specific prioritization, although validation across a larger and more diverse panel of cell lines will be essential to establish generalizability.

Predictors based on different types of drug information showed modest agreement in how they ranked the same compounds (pairwise Spearman ρ = 0.16–0.40, **Fig. 5f**). To assess how each information family contributed to hit recovery, we removed one family at a time and repeated predictor selection. Removing chemical fingerprints or transcriptomic profiles produced the largest decreases in mean EF@5%, by 0.67 and 0.60, respectively (paired compound-bootstrap *P* = 0.0368 and 0.0131). Enrichment decreased in nine of ten splits after fingerprint removal and in all ten after transcriptomic-profile removal (additional paired Wilcoxon signed-rank tests, *P* = 0.00586 and 0.00195, respectively, **Fig. 5g**). At the 5% screening budget, each method selected 27 compounds per cell line in each test split. Across four cell lines and ten splits, the ensemble recovered an average of 8.68 measured hits per 27 compounds, compared with 4.25–6.03 for the six single-family predictors and 1.55 expected from random selection (**Fig. 5h**). The ensemble recovered 2.65 more hits per 27 compounds than the best single-family predictor, which used LLM-derived annotations. Together, these findings suggest that integrating heterogeneous drug representations improves hit recovery by leveraging their complementary predictive contributions.

### External validation identifies compounds with preferential PGCC suppression

We applied the trained ensemble to rank 21,367 LINCS and PRISM library entries not included in the original PGCC screen and experimentally evaluated six nominated compounds.^50,51^ Five of the six candidates had a maximum ECFP6 Tanimoto similarity below 0.70 to any compound in the original screen; deslanoside was the exception, sharing a similarity of 0.769 with digoxin (**Fig. 6a**). These results demonstrate that the ensemble could prioritize PGCC-suppressing candidates that were not simply close structural analogs of compounds in the original screening library.

**Figure 6.**
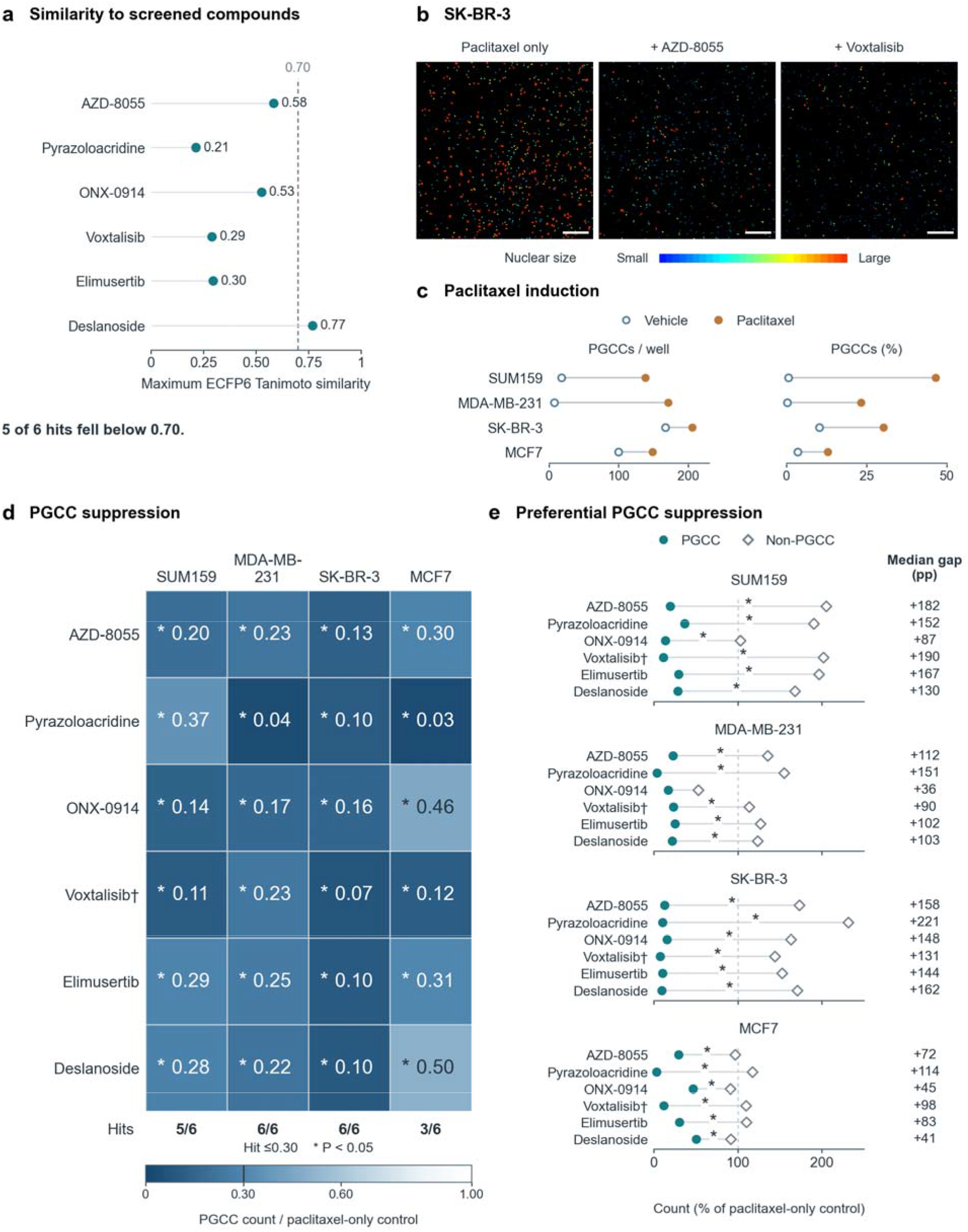
Paclitaxel-treated cultures. Preferential PGCC suppression in paclitaxel-treated cultures for validated compounds. a, Maximum ECFP6 Tanimoto similarity between each of the six confirmed hits and its nearest neighbor in the screened library. The dashed line marks 0.70. b, Full-field nuclear-size maps of SK-BR-3 cells treated with paclitaxel alone or with AZD-8055 or voxtalisib. Blue to red indicates increasing nuclear size on a common scale. Treatment images show the median PGCC-count well among three technical wells. Scale bars, 500 µm. c, Paired dots compare vehicle (open blue) and paclitaxel-only (filled orange) controls for mean PGCC count per well (left) and percentage of all cells classified as PGCCs (right). Each point is the median across three plates. d, Median PGCC count ratios for the six tested compounds in paclitaxel-treated cultures, normalized to the mean of same-plate paclitaxel-only control wells. Numbers show fold change. Darker blue indicates a lower PGCC count ratio on a continuous 0–1 scale. Hits have ratios ≤0.30 (≥70% reduction). Asterisks mark P < 0.05 from two-sided Welch’s t-tests comparing PGCC counts between drug-plus-paclitaxel and matched paclitaxel-only control wells. e, Median PGCC (filled circle) and non-PGCC (open diamond) counts as percentages of their respective same-plate paclitaxel-only control means (dashed line, 100%). The median gap is calculated from the within-well differences (normalized non-PGCC minus PGCC count) in percentage points. Positive values indicate a lower normalized PGCC count than non-PGCC count in the same well. Asterisks mark P < 0.05 from two-sided Welch’s t-tests comparing within-well log2(PGCC/non-PGCC) count ratios between treatment and matched control wells. Tests in d and e used three technical wells per treatment and 12 same-plate control wells. †Voxtalisib was tested at 0.5 µM. Other compounds were tested at 10 µM and paclitaxel at 1 µM.

We measured each candidate alone and alongside paclitaxel to assess PGCC suppression in untreated and chemotherapy-exposed cultures. Full-field nuclear-size maps illustrate the paclitaxel control and its combinations with AZD-8055 or voxtalisib (**Fig. 6b**). Paclitaxel increased the PGCC fraction in all four cell lines, with the clearest increases in absolute PGCC counts in SUM159 and MDA-MB-231 (**Fig. 6c**). Responses are medians of three replicates; complete dose and timing details are given in Methods. To measure the added effect of each candidate, we evaluated the combination of treatments relative to same-plate paclitaxel-only controls, using separate PGCC and non-PGCC counts.

All six compounds reduced PGCC counts by at least 70% in at least three paclitaxel-treated cell lines: AZD-8055, pyrazoloacridine, ONX-0914, voxtalisib, elimusertib and deslanoside. Five met the hit criteria in SUM159, six in both MDA-MB-231 and SK-BR-3, and three in MCF7 (**Fig. 6d**; **Supplementary Table 2**). AZD-8055 and voxtalisib met this threshold in all four lines. Across the 24 compound–cell-line combinations, comprising six compounds tested in four cell lines, PGCC counts were lower than paclitaxel only controls (two-sided Welch’s t-tests, all P < 0.001). For the six hits, we calculated selectivity in each well by subtracting the normalized PGCC count from the normalized non-PGCC count, each expressed as a percentage of its paclitaxel-only control. All 24 compound–cell-line pairs had positive selectivity scores, with a median of 122.3 percentage points across pairs (range, 35.7–221.1) (**Fig. 6e**). Within-well log2(PGCC/non-PGCC) count ratios were lower than those in matched controls in all 24 comparisons, with 23 reaching P < 0.05 (two-sided Welch’s t-tests; **Fig. 6e**; **Supplementary Data 1**). These paired measurements indicate that the candidates preferentially reduce PGCC abundance relative to non-PGCC populations, rather than simply causing generalized cytotoxicity. Six-drug response measurements and prediction–measurement agreement under both treatment conditions are provided in **Supplementary Tables 2–3**. These results provide proof-of-concept validation of model-guided discovery in a previously untested compound library, identifying novel PGCC-suppressing candidates with preferential effects on PGCCs over non-PGCC populations.

## Discussion

The present framework brings measured morphological and transcriptional responses and PGCC-guided feature selection into prediction across four breast cancer cell lines of different molecular subtypes. At the same testing budget, the multimodal ensemble recovered 5.6 times as many hits as random selection and 2.5 times as many as the best individual predictor. Experimental testing of nominated external compounds then identified six with preferential PGCC suppression in paclitaxel-treated cultures. Together, these findings establish a scalable, data-driven strategy to extend PGCC-directed drug discovery beyond experimentally screened compounds and accelerate the identification of candidate therapeutics.

Effective drug-response prediction requires identifying and preserving biologically meaningful information within high-dimensional datasets. Our feature-selection results demonstrate that this can be achieved by integrating biological knowledge into the selection process, retaining predictive information in a smaller set of measurements. The selected Cell Painting features captured variation in nuclear shape, nuclear-edge DNA intensity, and the spatial organization of RNA and endoplasmic-reticulum staining (**Fig. 3a, b**). ^9,11,28^ These findings suggest that PGCC-associated drug responses are encoded not simply by cell enlargement, but also by changes in nuclear and cytoplasmic organization. A similar principle applied to gene expression: biological prioritization reduced the 3,407 measured CIGS features to smaller gene sets, including 481 LLM-prioritized features (85.9% reduction). Enrichment of cell-cycle pathways and higher average predictive performance than size-matched random gene lists further support the ability of biological knowledge to identify informative expression features (**Fig. 3d, e**). The LLM ratings also grouped drugs with known links to polyploidization. The cluster with the highest ratings across six cell-cycle processes included all 16 drugs annotated for microtubules or tubulin and 18 of the 19 annotated for Aurora kinases (**Fig. 3f**). These results establish biologically informed feature selection as an effective strategy for distilling complex morphological and molecular measurements into compact, interpretable representations of drug response while retaining information relevant to PGCC biology.

Beyond feature selection, these results highlight that the predictive value of biological information depends on how it is represented and combined. Auto-encoded fingerprints and contrastively encoded target and pathway annotations each gave higher held-out response correlations than their raw features (**Fig. 4g**). These results suggest that representation learning can make existing drug information more useful for predicting PGCC responses. With TabPFN combining the predictions, MLP base predictors gave higher held-out correlations than HGB, RF, or SVM in all ten matched splits (**Fig. 4i**). At the 5% testing budget, the MLP ensemble also achieved greater hit enrichment than the SVM ensemble (5.46-fold versus 3.64-fold). Integrating multiple information families provided a substantial advantage in compound prioritization over individual-family predictors. Their compound rankings showed partial agreement (**Fig. 5f**). At the same testing budget, the ensemble recovered more measured hits on average than any of the six single-family predictors (**Fig. 5h**). These findings highlight the importance of integrating partially overlapping but functionally complementary information sources, consistent with observations across other biological assay systems.^34^ Removing chemical fingerprints produced the largest observed decrease in hit enrichment (**Fig. 5g**). Five of the six validated hits fell below the 0.70 similarity threshold for a close structural neighbor in the original screen. The ensemble therefore nominated active compounds beyond close structural neighbors of the screening library.

The predicted and validated compounds included agents with established anticancer activity as well as candidates whose effects on PGCCs have not been previously reported. Notably, AZD-8055 and voxtalisib, both inhibitors of mTOR signaling, met the PGCC suppression threshold across all four paclitaxel-treated cell lines, consistent with previous evidence implicating mTOR in dormant PGCC formation.^52,53,54^ AZD-8055 has also been shown to prevent BMS-777607-induced polyploidy and restore sensitivity to subsequent chemotherapy, including paclitaxel, in breast cancer models.^55^ Other validated compounds have been investigated in breast cancer or broader oncology settings, including voxtalisib in combination with letrozole,^56^ pyrazoloacridine as a single agent in metastatic breast cancer,^57^ elimusertib in DNA-damage-response-deficient tumors and in combination with cisplatin,^58^ ^59^ and ONX-0914 in breast cancer cell models.^60^ Evidence from other cancer settings provides further pharmacological context for evaluating these compounds against breast cancer PGCCs. For example, combination studies have examined voxtalisib with temozolomide in high-grade glioma ^61^ and pyrazoloacridine with carboplatin in advanced cancer.^62^ Deslanoside, another validated candidate, has demonstrated preclinical activity in prostate cancer cells and xenografts ^63^ and enhanced arsenic-trioxide activity in lymphoma models.^64^ Importantly, although these compounds have prior anticancer evidence, their preferential activity against PGCCs during paclitaxel exposure has not been previously established. Thus, our findings reveal previously unrecognized anti-PGCC activity among existing therapeutic compounds. At the same time, the use of LLM-derived annotations and biologically guided feature selection in our prediction framework may favor compounds associated with better-characterized targets and mechanisms. This knowledge-guided approach offers a translational advantage by prioritizing compounds with existing pharmacological, preclinical, or clinical evidence, thereby providing opportunities for drug repurposing and potentially accelerating further development. However, it may be less effective at identifying compounds with entirely unexplored mechanisms or limited prior biological annotation. Overall, these findings highlight the potential of our framework to uncover new therapeutic applications for existing compounds while also identifying the need for complementary strategies to discover less-characterized vulnerabilities of PGCCs.

More broadly, this framework illustrates the potential of phenotype-specific prediction to uncover drug activities that conventional population-level measurements, such as overall cell viability, may overlook. By modeling responses of a defined cellular population, the approach could be extended beyond PGCCs to other phenotypic changes and transitions between cellular states, provided that corresponding measurements are available for model training. The demonstrated reuse of imaging profiles across biological assays further supports the potential for adapting this strategy to diverse phenotypic screening applications.^33^ Importantly, coupling predictive modeling with automated liquid handling and high-content imaging could transform this framework from a prediction tool into an adaptive discovery platform. Through iterative active learning, experimental results from each screening round would update the models and guide subsequent experiments by balancing the identification of promising compounds with the acquisition of information in underexplored chemical and cellular settings. Such a feedback-driven strategy could progressively improve screening efficiency, reduce redundant testing, and direct experimental resources toward the most informative and promising candidates.

An important next step is to establish the generalizability and therapeutic relevance of preferential PGCC suppression. The limited number of cell lines and sparse nature of breast cancer subtypes used in this study may have limited the benefits of cellular context. Future work should evaluate broader cell-line panels and explore alternative ways to represent cellular context. Our bootstrap intervals quantify variation from sampling compounds with fixed models and gene lists; they do not capture variation from retraining or gene-list selection. Although all six compounds met the PGCC suppression criterion in at least three of the four paclitaxel-treated cell lines, testing in patient-derived models is needed to assess their activity across clinically relevant tumor heterogeneity. Varying treatment schedules and duration will also distinguish inhibition of PGCC formation from suppression of established PGCCs, while post-treatment follow-up will determine whether these effects reduce tumor-cell regrowth. Finally, mechanistic and *in vivo* studies assessing drug exposure, tolerability, and antitumor efficacy will clarify the underlying vulnerabilities and translational potential of these compounds. Together, these studies will establish the broader applicability and therapeutic utility of the framework.

## Methods

### Screening data and response endpoints

Our published screen used the 2,726-compound APExBIO DiscoveryProbe L1052 library in SUM159, MDA-MB-231, MCF7 and SK-BR-3 cells. After excluding autofluorescent compounds that interfered with the imaging pipeline and removing duplicate entries, 2,691 compounds remained for response modeling.^18^ Cells were cultured for 24 h before 48 h of treatment with each compound alone at 10 µM. Live/dead and nuclear staining distinguished viable PGCC and non-PGCC populations, with PGCC classification based on nuclear size. PGCC and non-PGCC counts were normalized to plate-matched negative controls and log2-transformed. There were 2,690 evaluable compounds in SUM159 and MCF7 and 2,691 in MDA-MB-231 and SK-BR-3.

### Drug information and numerical representations

Chemical fingerprints comprised MACCS, PubChem and ECFP6 descriptors.^26,65^ Curated annotations came from APExBIO DiscoveryProbe L1052 vendor metadata. CIGS and LINCS supplied expression profiles,^29,50^ with CIGS gene sets restricted using PGCC-associated RNA analyses or LLM-guided prioritization. JUMP-CP supplied external morphology profiles.^28^ Replicate profiles were averaged within each of ten mechanism-guided reference compounds, followed by an equal-weight average across compounds. DMSO means were subtracted, and the 200 measurements with the largest absolute shifts were retained. Text and molecular embeddings were constructed from compound descriptions and SMILES, respectively.^26,31^ Data sources and feature lists are listed in **Supplementary Data 1**; processing procedures are given in **Supplementary Computational Methods**.

GPT-4o queries through its application programming interface (API) supplied binary target/pathway assignments and separate scores for drug associations with genes, pathways and six biological questions. Higher scores represented stronger rated mechanistic links. For LLM-prioritized genes, model inputs instead contained measured expression values.

Fingerprint autoencoders produced 64-dimensional vectors,^37^ while contrastive encoders produced 128-dimensional vectors from independently perturbed versions of the same input.^39^ We fitted these transforms to the drug features and reused them across supervised cross-validation folds. Variational autoencoders were fitted separately within each supervised training fold.^38^ Input-specific architectures and training settings are recorded in **Supplementary Data 1**. Baseline mutation, copy-number, CRISPR-dependency and RNA-seq profiles from DepMap and Cell Model Passports provided cellular context for each cell line.^66,67^ These profiles were reduced in dimension and combined with the drug inputs. Encoder fitting, cellular-context processing, and ensemble width-sensitivity protocols (**Fig. 4h** and **Supplementary Fig. 2**) are detailed in **Supplementary Computational Methods**.

### Individual predictors and ensemble selection

Individual predictors used MLP, RF, HGB, or SVM regression.^68–71^ The MLP used shared hidden layers and four cell-line-specific output layers. Each compound–cell-line input used its corresponding output layer to predict PGCC and non-PGCC responses. MLPs minimized mean squared error with equal weights across training records and the PGCC and non-PGCC outputs. Input-specific architectures and training settings are recorded in **Supplementary Data 1**. Base out-of-fold (OOF) predictions were generated by ten repetitions of ten-fold cross-validation, excluding each predicted compound from base-model fitting. Medians across repetitions supplied one value per compound, cell line and candidate predictor. This matrix was reused across ensemble splits, so base models generating ensemble-training inputs could have used responses from compounds later assigned to testing. Evaluation therefore held compounds out at the ensemble stage. We refitted the base models with and without ensemble-test compounds while keeping the selected inputs fixed (**Supplementary Table 1**).

Ensembles were fitted separately for each base algorithm. The comparison across MLP, RF, HGB, and SVM used 82 candidate input identifiers shared across all four algorithms, with each algorithm retaining its recorded preprocessing settings. For MLP screening, candidate pools reflected the availability of additional target and pathway annotations. Four cell-line indicators were appended to ensemble inputs. Candidate definitions and preprocessing assignments are given in **Supplementary Computational Methods** and **Supplementary Data 1**.

The five learned combination methods were TabPFN, CatBoost, XGBoost, LightGBM and two-layer stacking. Each was evaluated on the same ten repeated 80:20 train–test splits of compounds (outer splits).^46–48^ Splits were stratified by mean compound response, with all cell-line records for each compound kept together. Within five-fold inner cross-validation, candidate filtering used training correlations and predefined redundancy constraints. Eligible family combinations were selected by mean validation of Pearson correlation across cell lines. Predictor filtering was repeated on the full outer-training set before fitting the selected ensemble. Missing predictions were imputed from training-partition column medians. Detailed selection rules, imputation procedures, and combination-model settings are given in **Supplementary Computational Methods**.

The best individual predictor was selected using Pearson correlation pooled across outer-training rows, with test predictions taken from the corresponding base OOF column. The simple-average comparator averaged the log2 prediction columns selected by TabPFN in each split, using the same training-median imputation but excluding cell-line indicators.

### Performance measures and statistical comparisons

Pearson correlation and EF@5% were calculated separately for each cell line. A measured hit had a PGCC count ratio ≤0.30. Compounds were ranked by increasing predicted response, and the top 5% count was rounded upward to 27 compounds in test sets of 537–538. Cell-line metrics were averaged equally within each split, followed by the mean and sample standard deviation (SD) across ten overlapping splits. Hit-recovery curves, precision–recall curves and predicted-rank groups used the same matched predictions.

For N test compounds, let H be the number of measured hits, k = 0.05N the number selected, and h the hits among those k compounds. Enrichment and hit recovery are defined as:

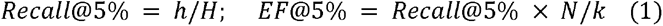

Algorithm summaries for **Fig. 4a–f** were descriptive; feature-set ranking and pooling procedures are detailed in **Supplementary Computational Methods**. For the ensemble comparison (**Fig. 4i**) and TabPFN-versus-averaging comparisons (**Tables 1–2**), we used two-sided overlap-corrected paired t-tests on ten matched four-line mean differences in Pearson correlation and EF@5%.^72^ For **Figs. 4h and 5h**, mean hit counts and the descriptive standard error of the mean (SEM) were calculated from ten split-level means across four cell lines. Feature-selection, individual-predictor cellular-context, and **Figure 5** comparisons used paired compound-bootstrap tests with pointwise 95% confidence intervals. Each draw preserved compound multiplicities across compared methods, available cell-line records and overlapping splits or repetitions, conditional on the fitted models. For family removal (**Fig. 5g**), additional two-sided paired Wilcoxon signed-rank tests assessed the ten split-level differences in four-line mean EF@5%. Representation-specific tests, resampling procedures, interval definitions, and SEM calculations are provided in **Supplementary Computational Methods**.

Feature-selection comparisons used matched compounds and ten repetitions of ten-fold compound-grouped cross-validation, with preprocessing and early stopping fitted on training compounds. Cell Painting comparisons evaluated the top 50, 100, 200, 300 or 500 features from the same reference-drug ranking (**Fig. 3c**). For **Fig. 3e**, each selected gene list was compared with five fixed size-matched random lists drawn from the same measured CIGS genes. All gene lists used 128-dimensional contrastive encoders fitted within the training folds and matched MLP settings. Reactome enrichment used one-sided hypergeometric tests with Benjamini–Hochberg correction across lists and pathways.^45,73^ Full protocols and numerical results are provided in **Supplementary Computational Methods** and **Supplementary Data 1**.

### Information-family contributions and cell-line comparisons

Each information family was removed in turn from the full 91-candidate pool, and predictor filtering and inner family selection were repeated within each outer-training partition. Comparisons used the same fitted base predictors and original test compounds. For ranking agreement, each retained family contributed the predictor with the highest training Pearson correlation averaged across cell lines. Pairwise Spearman correlations compared predicted rankings on jointly observed test compounds. For **Fig. 5h**, the same training criterion selected one predictor per split from each of the six families among all 91 candidates. Each method nominated 27 compounds from the same held-out pool per cell line and split. Detailed selection and averaging procedures are given in **Supplementary Computational Methods**.

For compounds measured in all four lines, we counted the lines meeting the hit threshold. On held-out compounds, we evaluated whether models predicted which of two lines had the lower PGCC count ratio, among pairs of measured responses differing by at least twofold. The cell-line mean baseline used each line’s mean log2 response to training compounds. Comparison eligibility, tie handling, and averaging are specified in **Supplementary Computational Methods**. Additional cellular-context comparisons are reported in **Supplementary Table 4**.

### External-library prediction and candidate characterization

The ensemble used additional LLM-derived binary target and pathway indicators when available, with separately selected inputs and fitted TabPFN configurations. Screened and library compounds used the same saved encoders and feature order. The external screening set comprised 21,367 LINCS and PRISM entries with at least one selected base prediction and no name or standardized parent-structure match to the PGCC screen. Missing selected base predictions were imputed using the medians of the corresponding training OOF predictions. Entries were ranked by increasing predicted PGCC responses within each cell line. Input availability, configuration details, and structure matching are described in **Supplementary Computational Methods**.

For each of the six confirmed hits, chemical resemblance was the maximum Tanimoto similarity of its standardized-structure ECFP6 fingerprint to the screened reference collection (**Fig. 6a**). We used a maximum similarity of 0.70 as a descriptive threshold for a close neighbor. Structure processing and candidate identities are described in **Supplementary Computational Methods**.

### Experimental assessment of nominated compounds

The six validation compounds were selected from the ensemble-ranked external library and tested in SUM159, MDA-MB-231, SK-BR-3 and MCF7 cells, alone and with paclitaxel. Cells attached for 24 hours before treatment. Responses were evaluated at 10 µM compounds, except voxtalisib at 0.5 µM. Paclitaxel (1 µM) and the test compound were added simultaneously. After 48 h of continuous exposure, images were analyzed with the original screening pipeline. Each treatment comprised three same-day technical wells. Counts for each population were divided by the mean of 12 corresponding same-plate vehicle or paclitaxel-only control wells and summarized by their median. The hit criterion was a PGCC count ratio of ≤0.30. Preferential suppression was quantified within each well as normalized non-PGCC minus PGCC counts, expressed in percentage points, and summarized by the median across technical wells (**Fig. 6e**). For **Fig. 6d**, two-sided Welch’s t-tests compared PGCC counts between three drug-plus-paclitaxel wells and 12 same-plate paclitaxel-only control wells. For **Fig. 6e**, the same test compared within-well log2(PGCC/non-PGCC) count ratios between these groups, retaining the pairing of the two populations within each well. Additional doses, controls, and prediction– measurement correlations are described in **Supplementary Experimental Methods** and **Supplementary Tables 2–3**. Individual-well data and statistical results are provided in **Supplementary Data 1**.

## Data availability

Data supporting the analyses are provided in **Supplementary Data 1** and the **Supplementary Information**. Public datasets used in this study are identified in the Methods and **Supplementary Information**. Additional processed data, trained models, and prediction files will be made publicly available through Zenodo upon journal publication.

## Code availability

The code used for feature preparation, model training, ensemble evaluation, statistical analysis, and figure generation will be made publicly available on GitHub upon journal publication.

## Acknowledgments and funding

This study was generously funded by start-up support from the UPMC Hillman Cancer Center, and the US National Institutes of Health grants P30CA047904, P50CA272218, R35GM150509, R21CA293424, R21CA302562, and R21EY039420. This research was supported in part by the University of Pittsburgh Center for Research Computing (NIH S10OD028483), through the resources provided.

## Author contributions

H.Y., C.-H.S., and Y.M. implemented and evaluated the prediction models and drug-target/pathway annotation datasets. H.-C.W. developed the Cell Painting features and contributed to LLM-derived annotations and associated methods. G.W. contributed to ensemble evaluation and methodology. O.B. performed chemical similarity and cross-cell-line ranking analyses. P.-R.L., H.-C.C., Y.Z. and J.C. contributed to the screening datasets and experimental validation of candidate hits. Y.-C.C. and Y.-C.C. supervised the study. H.Y., H.-C.W., G.W., Y.-C.C., and Y.-C.C. drafted the manuscript. All authors discussed the results, provided comments on the manuscript, and approved the final version.

## Competing interests

The authors declare no competing interests.

## Notes

### Competing Interest Statement

The authors have declared no competing interest.

## References

1. Bray, F., et al. Global cancer statistics 2022: GLOBOCAN estimates of incidence and mortality worldwide for 36 cancers in 185 countries. CA Cancer J. Clin. 74, 229–263 (2024). 10.3322/caac.21834.

2. Vasan, N., Baselga, J. & Hyman, D. M. A view on drug resistance in cancer. Nature 575, 299– 309 (2019). 10.1038/s41586-019-1730-1.

3. Carter, S. L. et al. Absolute quantification of somatic DNA alterations in human cancer. Nat. Biotechnol. 30, 413–421 (2012). 10.1038/nbt.2203.

4. Quinton, R. J. et al. Whole-genome doubling confers unique genetic vulnerabilities on tumour cells. Nature 590, 492–497 (2021). 10.1038/s41586-020-03133-3.

5. Bielski, C. M. et al. Genome doubling shapes the evolution and prognosis of advanced cancers. Nat. Genet. 50, 1189–1195 (2018). 10.1038/s41588-018-0165-1.

6. Gerashchenko, B. I. et al. Disentangling the aneuploidy and senescence paradoxes: a study of triploid breast cancers non-responsive to neoadjuvant therapy. Histochem. Cell Biol. 145, 497–508 (2016). 10.1007/s00418-016-1415-x.

7. Fei, F. et al. The number of polyploid giant cancer cells and epithelial-mesenchymal transition-related proteins are associated with invasion and metastasis in human breast cancer. J. Exp. Clin. Cancer Res. 34, 158 (2015). 10.1186/s13046-015-0277-8.

8. Amend, S. R. et al. Polyploid giant cancer cells: Unrecognized actuators of tumorigenesis, metastasis, and resistance. Prostate 79, 1489–1497 (2019). 10.1002/pros.23877.

9. Pienta, K. J., Hammarlund, E. U., Brown, J. S., Amend, S. R. & Axelrod, R. M. Cancer recurrence and lethality are enabled by enhanced survival and reversible cell cycle arrest of polyaneuploid cells. *Proc. Natl Acad. Sci. USA* 118, e2020838118 (2021). 10.1073/pnas.2020838118.

10. Saini, G. et al. Polyploid giant cancer cell characterization: new frontiers in predicting response to chemotherapy in breast cancer. Semin. Cancer Biol. 81, 220–231 (2022). 10.1016/j.semcancer.2021.03.017.

11. Schmidt, M. J. et al. Polyploid cancer cells reveal signatures of chemotherapy resistance. Oncogene 44, 439–449 (2025). 10.1038/s41388-024-03212-z.

12. Zhang, S. et al. Generation of cancer stem-like cells through the formation of polyploid giant cancer cells. Oncogene 33, 116–128 (2014). 10.1038/onc.2013.96.

13. Niu, N. et al. Linking genomic reorganization to tumor initiation via the giant cell cycle. Oncogenesis 5, e281 (2016). 10.1038/oncsis.2016.75.

14. Song, Y., Zhao, Y., Deng, Z., Zhao, R. & Huang, Q. Stress-induced polyploid giant cancer cells: unique way of formation and non-negligible characteristics. Front. Oncol. 11, 724781 (2021). 10.3389/fonc.2021.724781.

15. Sparano, J. A. et al. Weekly paclitaxel in the adjuvant treatment of breast cancer. N. Engl. J. Med. 358, 1663–1671 (2008). 10.1056/NEJMoa0707056.

16. Zhang, S., Mercado-Uribe, I. & Liu, J. Tumor stroma and differentiated cancer cells can be originated directly from polyploid giant cancer cells induced by paclitaxel. Int. J. Cancer 134, 508–518 (2014). 10.1002/ijc.28319.

17. Adan, A., Kiraz, Y. & Baran, Y. Cell proliferation and cytotoxicity assays. Curr. Pharm. Biotechnol. 17, 1213–1221 (2016). 10.2174/1389201017666160808160513.

18. Chen, H. C. et al. High-throughput drug screening for targeting polyploid cancer cells with an interactive web portal. Cancer Lett. 642, 218289 (2026). 10.1016/j.canlet.2026.218289.

19. Iorio, F. et al. A landscape of pharmacogenomic interactions in cancer. Cell 166, 740–754 (2016). 10.1016/j.cell.2016.06.017.

20. Sharifi-Noghabi, H., Zolotareva, O., Collins, C. C. & Ester, M. MOLI: multi-omics late integration with deep neural networks for drug response prediction. Bioinformatics 35, i501– i509 (2019). 10.1093/bioinformatics/btz318.

21. Liu, Q., Hu, Z., Jiang, R. & Zhou, M. DeepCDR: a hybrid graph convolutional network for predicting cancer drug response. Bioinformatics 36, i911–i918 (2020). 10.1093/bioinformatics/btaa822.

22. Park, S., Soh, J. & Lee, H. Super. FELT: supervised feature extraction learning using triplet loss for drug response prediction with multi-omics data. BMC Bioinformatics 22, 269 (2021). 10.1186/s12859-021-04146-z.

23. Chen, J. & Zhang, L. A survey and systematic assessment of computational methods for drug response prediction. Brief. Bioinform. 22, 232–246 (2021). 10.1093/bib/bbz164.

24. Xia, F. et al. A cross-study analysis of drug response prediction in cancer cell lines. Brief. Bioinform. 23, bbab356 (2022). 10.1093/bib/bbab356.

25. Bernett, J. et al. Critical evaluation of drug response prediction models with DrEval. Nat. Commun. 17, 4238 (2026). 10.1038/s41467-026-72903-w.

26. Ma, Y. et al. High-throughput empirical and virtual screening to discover novel inhibitors of polyploid giant cancer cells in breast cancer. Anal. Chem. 97, 5498–5506 (2025). 10.1021/acs.analchem.4c05138.

27. Seal, S. et al. Cell Painting: a decade of discovery and innovation in cellular imaging. Nat. Methods 22, 254–268 (2025). 10.1038/s41592-024-02528-8.

28. Chandrasekaran, S. N. et al. Three million images and morphological profiles of cells treated with matched chemical and genetic perturbations. Nat. Methods 21, 1114–1121 (2024). 10.1038/s41592-024-02241-6.

29. Xiang, L. et al. High-throughput profiling of chemical-induced gene expression across 93,644 perturbations. Nat. Methods 22, 1954–1963 (2025). 10.1038/s41592-025-02781-5.

30. Knox, C. et al. DrugBank 6.0: the DrugBank Knowledgebase for 2024. Nucleic Acids Res. 52, D1265–D1275 (2024). 10.1093/nar/gkad976.

31. Zheng, Y. et al. Large language models for scientific discovery in molecular property prediction. *Nat*. Mach. Intell. 7, 437–447 (2025). 10.1038/s42256-025-00994-z.

32. Carli, F. et al. Learning and actioning general principles of cancer cell drug sensitivity. Nat. Commun. 16, 1654 (2025). 10.1038/s41467-025-56827-5.

33. Simm, J. et al. Repurposing high-throughput image assays enables biological activity prediction for drug discovery. Cell Chem. Biol. 25, 611–618.e3 (2018). 10.1016/j.chembiol.2018.01.015.

34. Moshkov, N. et al. Predicting compound activity from phenotypic profiles and chemical structures. Nat. Commun. 14, 1967 (2023). 10.1038/s41467-023-37570-1.

35. Hauptmann, T. & Kramer, S. A fair experimental comparison of neural network architectures for latent representations of multi-omics for drug response prediction. BMC Bioinformatics 24, 45 (2023). 10.1186/s12859-023-05166-7.

36. Neve, R. M. et al. A collection of breast cancer cell lines for the study of functionally distinct cancer subtypes. Cancer Cell 10, 515–527 (2006). 10.1016/j.ccr.2006.10.008.

37. Hinton, G. E. & Salakhutdinov, R. R. Reducing the dimensionality of data with neural networks. Science 313, 504–507 (2006). 10.1126/science.1127647.

38. Kingma, D. P. & Welling, M. Auto-encoding variational Bayes. Preprint at 10.48550/arxiv.1312.6114 (2013).

39. Chen, T., Kornblith, S., Norouzi, M. & Hinton, G. A simple framework for contrastive learning of visual representations. Proc. Mach. Learn. Res. 119, 1597–1607 (2020). https://proceedings.mlr.press/v119/chen20j.html.

40. Fei, F. et al. The subcellular location of cyclin B1 and CDC25 associated with the formation of polyploid giant cancer cells and their clinicopathological significance. Lab. Invest. 99, 483–498 (2019). 10.1038/s41374-018-0157-x.

41. Zhou, M. et al. Single-cell morphological and transcriptome analysis unveil inhibitors of polyploid giant breast cancer cells in vitro. *Commun*. Biol. 6, 1301 (2023). 10.1038/s42003-023-05674-5.

42. Tagal, V. & Roth, M. G. Loss of Aurora kinase signaling allows lung cancer cells to adopt endoreplication and form polyploid giant cancer cells that resist antimitotic drugs. Cancer Res. 81, 400–413 (2021). 10.1158/0008-5472.CAN-20-1693.

43. Chen, J.-G., Yang, C.-P. H., Cammer, M. & Horwitz, S. B. Gene expression and mitotic exit induced by microtubule-stabilizing drugs. Cancer Res. 63, 7891–7899 (2003).

44. Zhang, X. et al. Targeting polyploid giant cancer cells potentiates a therapeutic response and overcomes resistance to PARP inhibitors in ovarian cancer. Sci. Adv. 9, eadf7195 (2023). 10.1126/sciadv.adf7195.

45. Milacic, M. et al. The Reactome Pathway Knowledgebase 2024. Nucleic Acids Res. 52, D672–D678 (2024). 10.1093/nar/gkad1025.

46. Hollmann, N. et al. Accurate predictions on small data with a tabular foundation model. Nature 637, 319–326 (2025). 10.1038/s41586-024-08328-6.

47. 47. Grinsztajn, L., et al. TabPFN-3: technical report. Preprint at 10.48550/arxiv.2605.13986 (2026).

48. Wolpert, D. H. Stacked generalization. Neural Netw. 5, 241–259 (1992). 10.1016/s0893-6080(05)80023-1.

49. Mnich, K., Kitlas Golińska, A., Polewko-Klim, A. & Rudnicki, W. R. Bootstrap bias corrected cross validation applied to super learning. In Computational Science – ICCS 2020. Lect. Notes Comput. Sci. 12139, 550–563 (2020). 10.1007/978-3-030-50420-5_41.

50. Subramanian, A. et al. A next generation Connectivity Map: L1000 platform and the first 1,000,000 profiles. Cell 171, 1437–1452.e17 (2017). 10.1016/j.cell.2017.10.049.

51. Corsello, S. M. et al. Discovering the anticancer potential of non-oncology drugs by systematic viability profiling. Nat. Cancer 1, 235–248 (2020). 10.1038/s43018-019-0018-6.

52. You, B. et al. AMPK-mTOR-mediated activation of autophagy promotes formation of dormant polyploid giant cancer cells. Cancer Res. 82, 846–858 (2022). 10.1158/0008-5472.CAN-21-2342.

53. Chresta, C. M., et al. AZD8055 is a potent, selective, and orally bioavailable ATP-competitive mammalian target of rapamycin kinase inhibitor with in vitro and in vivo antitumor activity. Cancer Res. 70, 288–298 (2010). 10.1158/0008-5472.CAN-09-1751.

54. Yu, P. et al. Characterization of the activity of the PI3K/mTOR inhibitor XL765 (SAR245409) in tumor models with diverse genetic alterations affecting the PI3K pathway. Mol. Cancer Ther. 13, 1078–1091 (2014). 10.1158/1535-7163.MCT-13-0709.

55. Sharma, S. et al. Prevention of BMS-777607-induced polyploidy/senescence by mTOR inhibitor AZD8055 sensitizes breast cancer cells to cytotoxic chemotherapeutics. Mol. Oncol. 8, 469–482 (2014). 10.1016/j.molonc.2013.12.014.

56. Blackwell, K. et al. Phase I/II dose-escalation study of PI3K inhibitors pilaralisib or voxtalisib in combination with letrozole in patients with hormone-receptor-positive and HER2-negative metastatic breast cancer refractory to a non-steroidal aromatase inhibitor. Breast Cancer Res. Treat. 154, 287–297 (2015). 10.1007/s10549-015-3615-9.

57. Ramaswamy, B., Mrozek, E., Kuebler, J. P., Bekaii-Saab, T. & Kraut, E. H. Phase II trial of pyrazoloacridine (NSC#366140) in patients with metastatic breast cancer. Invest. New Drugs 29, 347–351 (2011). 10.1007/s10637-009-9338-1.

58. Yap, T. A. et al. Phase Ib basket expansion trial and alternative-schedule dose-escalation study of ATR inhibitor elimusertib in advanced solid tumors with DNA damage response defects. Cancer Discov. 15, 2019–2035 (2025). 10.1158/2159-8290.cd-24-1500.

59. Parikh, M. et al. The ATR inhibitor elimusertib in combination with cisplatin in patients with advanced solid tumors: a California Cancer Consortium Phase I trial (NCI 10404). Cancer Res. Commun. 5, 1946–1951 (2025). 10.1158/2767-9764.CRC-25-0305.

60. Kondakova, I. et al. Association of proteasome activity and pool heterogeneity with markers determining the molecular subtypes of breast cancer. Cancers 17, 159 (2025). 10.3390/cancers17010159.

61. Wen, P. Y. et al. Phase I dose-escalation study of the PI3K/mTOR inhibitor voxtalisib (SAR245409, XL765) plus temozolomide with or without radiotherapy in patients with high-grade glioma. Neuro Oncol. 17, 1275–1283 (2015). 10.1093/neuonc/nov083.

62. Adjei, A. A. et al. A phase I and pharmacologic study of pyrazoloacridine (NSC 366140) and carboplatin in patients with advanced cancer. Invest. New Drugs 20, 297–304 (2002). 10.1023/A:1016237426846.

63. Liu, M. et al. The cardiac glycoside deslanoside exerts anticancer activity in prostate cancer cells by modulating multiple signaling pathways. Cancers 13, 5809 (2021). 10.3390/cancers13225809.

64. Li, G. et al. Arsenic trioxide promotes a glue-like interaction to drive STUB1-mediated NPM-ALK degradation in ALK+ ALCL. Exp. Hematol. Oncol. 15, 66 (2026). 10.1186/s40164-026-00800-5.

65. Rogers, D. & Hahn, M. Extended-connectivity fingerprints. J. Chem. Inf. Model. 50, 742–754 (2010). 10.1021/ci100050t.

66. Arafeh, R., Shibue, T., Dempster, J. M., Hahn, W. C. & Vazquez, F. The present and future of the Cancer Dependency Map. Nat. Rev. Cancer 25, 59–73 (2025). 10.1038/s41568-024-00763-x.

67. van der Meer, D. et al. Cell Model Passports—a hub for clinical, genetic and functional datasets of preclinical cancer models. Nucleic Acids Res. 47, D923–D929 (2019). 10.1093/nar/gky872.

68. Breiman, L. Random forests. Mach. Learn. 45, 5–32 (2001). 10.1023/a:1010933404324.

69. Pedregosa, F. et al. Scikit-learn: machine learning in Python. J. Mach. Learn. Res. 12, 2825–2830 (2011). https://jmlr.org/papers/v12/pedregosa11a.html.

70. Smola, A. J. & Schölkopf, B. A tutorial on support vector regression. Stat. Comput. 14, 199– 222 (2004). 10.1023/b:stco.0000035301.49549.88.

71. Paszke, A. et al. PyTorch: an imperative style, high-performance deep learning library. Adv. Neural Inf. Process. Syst. 32 (2019). https://proceedings.neurips.cc/paper/2019/hash/bdbca288fee7f92f2bfa9f7012727740-Abstract.html.

72. Nadeau, C. & Bengio, Y. Inference for the generalization error. Mach. Learn. 52, 239–281 (2003). 10.1023/A:1024068626366.

73. Benjamini, Y. & Hochberg, Y. Controlling the false discovery rate: a practical and powerful approach to multiple testing. J. R. Stat. Soc. B 57, 289–300 (1995). 10.1111/j.2517-6161.1995.tb02031.x.

