## Supplementary Information for "Multimodal Virtual Screening Identifies Candidate Inhibitors of Polyploid Giant Cancer Cells Across Breast Cancer Subtypes"

PGCC drug response prediction supplementary material

**Supplementary Figures 1–2**, **Tables 1–4** and Methods accompany the manuscript. References are numbered independently. **Supplementary Data 1** contains feature lists, source information, model settings, statistical comparisons and validation measurements. **Prompt sets 1–5** contain the supplied scoring instructions.

### Supplementary Computational Methods

#### Fingerprint autoencoders

MACCS, PubChem and ECFP6 inputs contained 166, 881 and 1,024 binary features, respectively, and were supplied as 0/1 values. The first 2,181 rows in source order were used for reconstruction training, with the remaining 545 used for validation.

Fingerprint MLP-AEs used a fixed 64-dimensional bottleneck and were trained on screened fingerprints using mean squared reconstruction error. Their saved weights encoded both screened and external-library compounds. Screening input combinations are listed in **Supplementary Data 1**.

Each encoder comprised fully connected layers of 128, 64 and 64 units. Rectified linear activations followed the first two layers; the latent output was linear. The decoder used layers of 64 and 128 rectified linear units followed by a linear reconstruction layer. Training minimized mean squared reconstruction error with Adam, using learning rate 0.001, batch size 32, 100 epochs and seed 42. Final-epoch weights were retained.

The same fitted encoder and ordered input columns were used to transform screened and library compounds. Concatenating individual latent representations produced single-, paired- and three-fingerprint feature sets with 64, 128 and 192 features, respectively. These unsupervised encoders were shared before supervised cross-validation. Contrastive encoders operating on these representations retained their fitted imputation, scaling and feature order for subsequent transformation.

#### Contrastive encoders

Contrastive learning^1^ used a separate encoder for each drug-feature input, with a 512-unit rectified linear hidden layer, dropout of 0.10 and a linear 128-dimensional output. A projection head contained two 64-unit layers with an intervening rectified linear activation. Projection outputs were normalized to unit length for the contrastive loss. The downstream predictors used the unit-normalized 128-dimensional encoder outputs.

Missing feature entries were filled with column means calculated from available input rows, and each feature was standardized using the same rows. Each training input generated two independent views through 5% feature dropout and Gaussian noise with standard deviation 0.01. Dropped features were set to zero and retained features were divided by 0.95. The loss compared the two views of each row with other rows in the minibatch, using temperature 0.10. Training used AdamW for 60 epochs with batch size 256, learning rate 0.001, weight decay 0.0001, gradient-norm clipping at one and seed 42. The loss compared perturbed views within each input; no loss term aligned different feature inputs.

These encoders were fitted before the supervised folds and reused across them. The fitting inputs retained separate compound–cell-line rows, so a compound could contribute repeated feature vectors across cell lines even though cell-line context was excluded from the encoder. For fingerprint-AE inputs, contrastive fitting used the screened compounds and the same training settings. Saved feature order, imputation values, scaling parameters and encoder weights were reused for library transformation.

#### Variational autoencoders

For the MACCS, PubChem and ECFP6 VAE candidate predictors, the VAE^2^ used 64 latent dimensions, encoder hidden layers of 512 and 256 rectified linear units, and a decoder with the reversed hidden-layer sizes. Separate linear heads produced the latent mean and log variance. Training sampled the latent vector using the reparameterization rule; prediction used the latent mean.

Within each supervised fold, feature means and standard deviations were fitted on the training rows. The VAE was then trained on those standardized rows alone, using the assembled feature matrix supplied to the predictor, including appended cell-line context where enabled. Training used Adam with learning rate 0.0005, batch size 256 and 120 epochs. The objective was mean squared reconstruction error plus the mean Kullback–Leibler divergence from a standard-normal prior, with a fixed coefficient of one. Final-epoch VAE weights were retained. The same scaler and encoder transformed the held-out rows. Subsequent MLP training used its separately recorded predictor settings.

#### Cell-painting processing

The 200-feature and nuclei-only representations used the JUMP-CP compound_interpretable v1.0 release^3^ after median absolute deviation normalization and inverse-normal transformation. Compound profiles were selected by retaining the first source-order well for each JUMP compound identifier. The control vector was the feature-wise arithmetic mean of 384 DMSO wells from source_11, plate DMSO71-102_1. This vector was subtracted from each selected compound profile.

The predefined column sets comprised 200 features and 1,055 nuclei features, respectively. Their order was preserved across screened compounds and external-library inputs.

Feature selection used ten mechanism-guided reference compounds: five Aurora inhibitors (tozasertib, AMG-900, MK-5108, ZM 447439 and CCT129202), two PARP inhibitors (INO-1001 and AG-14361) and three microtubule-targeting drugs (mebendazole, epothilone B and ABT-751). Compound identifiers are listed in **Supplementary Data 1**. Compound identities were resolved by JUMP identifier. Replicate wells were averaged within each reference compound, followed by an equally weighted mean across the ten compounds. The feature-wise mean of 384 DMSO wells was subtracted, and measurements were ranked by the absolute shift to select the top 50, 100, 200, 300 or 500 features. The nuclei-only input retained all 1,055 nuclei measurements. Reference profiles were replicate-averaged for selection; model-input profiles followed the separate first-well rule described above. **The 200-column input retained 194 nontechnical measurements and six technical columns**.

##### RNA-guided CIGS gene selection

The SUM159 and MDA-MB-231 single-cell inputs were drawn from PGCC-versus-non-PGCC marker tables. The author-reported selection criteria were adjusted P < 0.05 and absolute average log2 fold change > 0.5. Intersection with measured CIGS genes yielded inputs of 19 and 29 genes, respectively.

The SUM159 bulk RNA input retained 1,614 CIGS-overlapping genes at source FDR < 0.05 and absolute signed linear fold change > 1. In the source table, signed fold change equals the PGCC/non-PGCC expression ratio when that ratio is at least one, and minus its reciprocal otherwise. The MDA-MB-231 up/down genes set matched the union of the retained upregulated and downregulated lists, comprising 37 and 41 CIGS-overlapping genes, respectively.

The SUM159 bulk + single-cell input combined genes from the single-cell marker analysis and the bulk PGCC-versus-non-PGCC contrast. Gene symbols were standardized and duplicate symbols retained the row with the lowest source P value. Genes present in either source were eligible; genes with opposite directions across the two sources were excluded. Signed normal scores derived from two-sided source P values were combined using the Stouffer method. Benjamini–Hochberg adjustment^4^ was applied across the surviving genes before CIGS intersection. Adjusted P < 0.05 and absolute mean stored effect > 1 yielded 1,556 genes.

The combined-input effect threshold used the mean of bulk signed linear fold changes and single-cell log2 fold changes. This quantity is a feature-selection score on mixed scales. The bulk + single-cell input was included in the candidate predictor comparisons.

##### LLM-prioritized genes and CIGS feature membership

The LLM-prioritized genes input contains 481 expression columns. The reported retention rule was mean gene rating ≥0.60 with at least two of six biological-question ratings ≥0.65. Exact membership intersections are calculated from the fixed gene lists.

##### Library target and pathway annotations

For the LINCS and PRISM library, target and pathway annotations were assembled from ChEMBL mechanism-of-action records, ChEMBL bioactivity records^5^ and DGIdb drug–gene interactions^6^. Target and pathway names were mapped to the training categories using UniProt^7^ and Reactome identifiers^8^. This library extension was separate from the APExBIO vendor annotations used for the screened compounds.

Compounds were linked to ChEMBL through PubChem identifiers. We retained all matched mechanism-of-action records. Bioactivity records were restricted to human binding assays with IC50, Ki, Kd or EC50 measurements and pChEMBL ≥6; the ten most potent targets were retained per compound. DGIdb interactions were queried by name and checked against ChEMBL identifiers, including parent-compound aliases. Targets were mapped through UniProt to Reactome pathways and assigned to 346 target and 30 pathway features in the training matrices through a curated dictionary.

##### Binary annotation schemas

Expanded training matrices contained 1,651 target and 525 pathway columns (2,176 combined). Additional-label subsets contained 1,317 target and 500 pathway columns (1,817 combined). Additional labels were LLM-derived assignments absent from the curated categories. Combined matrices used target/pathway prefixes and explicit parent labels for retained composite pathways. GPT-4o was queried for screened compounds and library entries with a ChEMBL identifier or curated target information. Missing or uncertain assignments were encoded as zero.

For binary target/pathway annotation, GPT-4o was queried once per compound at temperature 0 with JSON output, using compound name, CAS number and SMILES as inputs and the prompt reproduced in **S5**.

##### LLM-derived score tables

The complete tables contained 35 gene, 61 pathway and six mechanistic-question scores for the screened compounds. Higher ratings denoted stronger mechanistic links, not potency or effect direction. Each drug was queried once per gene, pathway or question. Gene/pathway labels were mapped as very high 1, high 0.75, medium 0.5, low 0.25 and unknown 0. The direct six-question baseline requested numeric scores from 0.00 to 1.00 with up to two decimal places (**S4**). The prompts and score mappings are reproduced below.

The six mechanistic ratings were aligned to screened compounds by rack/well identifier. The missing G1-arrest rating for TAK-285 was encoded as zero.

##### LLM4SD molecular embeddings

The LLM4SD input comprised 64 numeric molecular embedding features per compound, aligned to the response data by rack/well identifier.

#### Supplied gene, pathway and mechanism scoring prompts

The drug-level prompts below describe gene and pathway ratings (**S1–S2**), their label-to-score mapping (**S3**), direct six-question numeric scoring (**S4**), and binary target/pathway annotation (**S5**).

#### **Prompt set 1. Gene scoring prompts (35 genes)**

Each gene was scored in a separate run over all compounds. Within a run, the four messages below were sent in order, with the fourth message repeated once per compound.

##### **Prompt 1.1 System message**

You are an expert in cancer pharmacology, cell-cycle biology, and mechanism-of-action reasoning.

Your task is to judge how strongly a drug is mechanistically connected to this gene:

{TARGET}

Interpret the gene name as the gene or its encoded protein product.

Use the drug's known or widely accepted mechanism of action as the primary basis for scoring.

Scoring rule:

Score the relationship based on whether the drug would be expected to directly target, regulate, inhibit, activate, destabilize, or consistently alter the activity, abundance, or function of this gene/protein in cancer cells.

Important:

- Direct binding is not required.

- However, the relationship must be specific and mechanistically meaningful.

- Do NOT assign a high score just because the drug affects the same broad pathway or general cell-cycle biology.

- Broad downstream consequences alone should usually be scored low or medium, not high.

- Weak, generic, or speculative associations should be scored low or unknown.

Use these labels:

- very_high = drug directly targets this gene/protein, or is a well-known strong regulator with a specific and reliable mechanistic effect on it

- high = drug has a well-established and strong mechanistic effect on this gene/protein

- medium = drug has a plausible but indirect, context-dependent, or less central mechanistic connection

- low = only weak, broad, or peripheral connection

- unknown = no reliable relationship can be recalled

Evidence type must be exactly one of:

- direct_target

- moa_to_gene

- indirect_contextual_link

- unknown

Confidence should reflect how certain the mechanistic relationship is, from 0 to 1.

Do not hallucinate niche evidence.

Prefer established pharmacology over speculation.

Later, when given one drug, return exactly one JSON object only.

##### **Prompt 1.2 Output-format message (user)**

```
You will receive one drug at a time.

Return exactly one JSON object in this format:

{
    "drug_number": 1,
    "drug": "DrugName",
    "likelihood": "medium",
    "evidence_type": "moa_to_gene",
    "confidence": 0.64
}

Rules:
- Copy drug_number exactly as given
- Copy drug exactly as given
- Return exactly one JSON object
- No markdown
- No explanation outside JSON

Reply only with: yes
```

##### **Prompt 1.3 Confirmation message (user)**

Great. From now on, for each drug I send, estimate whether it influences the gene/protein '{TARGET}'. Return exactly one JSON object only. Use the exact input drug_number and exact input drug name. If unknown, use likelihood = 'unknown' and evidence_type = 'unknown'. If you understand, answer yes.

##### **Prompt 1.4 Per-compound scoring message (user)**

Gene of interest: {TARGET}

Using the drug's known mechanism of action, judge how strongly this drug is mechanistically connected to this gene/protein.

Focus on whether the drug would directly target, regulate, inhibit, activate, destabilize, or consistently alter this gene/protein in cancer cells.

Do not use broad pathway similarity alone.

Input:

drug_number: {DRUG_NUMBER}

drug: {DRUG_NAME}

Return exactly one JSON object with keys:

drug_number, drug, likelihood, evidence_type, confidence

#### **Prompt set 2. Pathway scoring prompts (61 pathways)**

The pathway prompts are identical to the gene prompts in **Prompt set 1** except for one line of the system message, shown below. Each pathway was scored in a separate run over all compounds.

##### **Prompt 2.1 System message, changed line**

Your task is to judge how strongly a drug is mechanistically connected to this pathway:

#### **Prompt set 3. Gene and pathway label-to-score mapping**

| **Label returned by the model** | **Stored score** |
| --- | --- |
| very_high | 1.00 |
| high | 0.75 |
| medium | 0.50 |
| low | 0.25 |
| unknown | 0.00 |

#### **Prompt set 4. Direct six-question scoring prompts**

Each mechanism was scored in a separate run over all compounds. Within a run, the four messages below were sent in order, with the fourth message repeated once per compound.

##### **Prompt 4.1 System message**

You are a biomedical research expert, with extensive knowledge of small molecules and cell biology.

##### **Prompt 4.2 Opening message (user)**

Do you know about how small molecules regulate {MECHANISM}?

##### **Prompt 4.3 Confirmation message (user)**

The task concerns a planned drug screen experiment. For a list of compound names, evaluate the likelihood of each compound regulating {MECHANISM}. Based on your knowledge, provide a regulatory likelihood score. Each score must be between 0.00 and 1.00: - 1.00 = strongest likelihood that the compound regulates this process - 0.00 = no effect or not related Return all results strictly between 0 and 1 (inclusive) with up to two decimal places. If you understand, please answer yes.

##### **Prompt 4.4 Per-compound scoring message (user)**

The compound name is "{DRUG_NAME}". Return your answer as a single JSON object ONLY, with no extra text, in this format: {"Drug Name": "<string>", "Likelihood": "<0.00-1.00>", "Reasoning": "<short explanation>"}

##### **Direct six-question scoring procedure**

Mechanism likelihood scores were obtained by querying GPT-4o once per compound for each of six PGCC-related processes: cell cycle, endoreduplication, mitotic slippage, cytokinesis failure, G1 arrest and G2/M checkpoint failure. The prompt requested a numeric score from 0.00 (no effect or not related) to 1.00 (strongest likelihood), with up to two decimal places, and a short explanation in JSON. Each mechanism was queried separately, with one compound per request at temperature 0.2 in a single pass.

#### **Prompt set 5. Expanded binary annotations**

##### **Prompt 5.1 System message**

```
You are a pharmacology and medicinal chemistry expert.

When given a drug's name (and optional identifiers like PubChem CID, InChIKey, ChEMBL ID, CAS number, or SMILES), produce a concise structured assessment of what is known about the compound's biological activity.
Return ONLY a JSON object with these exact fields:
{
  "is_known_drug": true/false,
  "identification_confidence": "high" | "medium" | "low" | "unknown",
  "primary_targets": ["TARGET_HGNC_SYMBOL_1", "TARGET_HGNC_SYMBOL_2"],
  "primary_pathways": ["PATHWAY_1", "PATHWAY_2"],
  "mechanism_summary": "1-2 sentences describing the mechanism of action",
  "therapeutic_class": "e.g. antibiotic, antineoplastic, antifungal, kinase inhibitor, research tool",
  "notes": "any caveats, alternative names, or reasons for low confidence"
}

Rules:
 - If the drug is unfamiliar, set is_known_drug to false and identification_confidence to "unknown".
   Do NOT hallucinate targets. Leave primary_targets and primary_pathways as empty arrays.
 - primary_targets must be HGNC-approved gene symbols (e.g. "EGFR", "JAK1") when known.
 - primary_pathways should be short broad categories (e.g. "MAPK Signaling", "PI3K/Akt/mTOR Signaling", "DNA Damage/DNA Repair").
 - Output JSON only, no surrounding text, no markdown code fences.
```

##### **Prompt 5.2 Per-compound message (user)**

Drug Name: {DRUG_NAME}

PubChem CID: {PUBCHEM_CID}

InChIKey: {INCHIKEY}

ChEMBL ID: {CHEMBL_ID}

CAS Number: {CAS_NUMBER}

SMILES: {SMILES}

What is known about this compound's mechanism, targets, and pathways?

Return JSON only.

Binary annotation procedure

Returned primary_targets and primary_pathways labels were encoded as binary assignments. Curated assignments were retained, and composite categories were split when their parts represented distinct concepts. Identical categories were merged within the curated schema. LLM-derived labels sharing a name with a curated category were stored separately with the suffix _1.

#### Cellular context in individual predictors

Cellular-context profiles were reduced using singular value decomposition or principal component analysis and combined with drug inputs together with source-availability indicators. Source datasets and feature definitions are listed in **Supplementary Data 1**. We compared individual MLP predictors with and without cellular context using one fixed drug-feature input from each family: ECFP6 fingerprints, combined curated target/pathway annotations, expanded LLM-derived binary target/pathway annotations, large text embeddings, the selected 200 Cell Painting measurements and expression values for LLM-prioritized genes. Ten repeated 80:20 compound splits (seeds 12–21) were shared across the comparisons. All cell-line records for each compound remained together. Within each input and split, the two arms used identical records, initialization and training settings; compound availability could differ between feature inputs.

The network retained shared hidden layers of 256, 128 and 64 units, dropout of 0.2 and four cell-line-specific output layers predicting PGCC and non-PGCC responses. The no-context arm set all standardized cellular-context coordinates to zero during both fitting and prediction, preserving the network structure and cell-line-specific outputs. Feature and response standardization used training rows only. Adam minimized unweighted mean squared error for 200 epochs with learning rate 0.001, weight decay 0.0001 and nominal batch size 512. Final-epoch weights supplied predictions, which were transformed back to the log2 count-ratio scale. Drug-feature lists were fixed, and no representation fitting or feature reselection was performed in this comparison.

Pearson correlation and EF@5% were calculated within each cell line and averaged equally across the four lines for each split. Context effects were paired differences (with minus without context). **Supplementary Fig. 1** shows mean differences with pointwise 95% basic compound-bootstrap intervals. Relative enrichment changes use the ratio of the mean EF@5% values across splits. Both arms retain cell-line-specific output layers. The paired compound-bootstrap procedure is specified below.

#### Cell-line identity comparison

Using the same six drug inputs, ten compound splits and training settings, we compared drug-only inputs, drug plus cell-line identity, and drug plus molecular context. All three conditions used one shared output layer predicting PGCC and non-PGCC responses. The drug-only condition set all context coordinates to zero. The identity condition placed training-standardized one-hot cell-line indicators in four of these coordinates and set the remainder to zero; the molecular-context condition retained all context coordinates. Input width, parameter count, initialization and training records were matched within each comparison. **Supplementary Table 4** reports mean EF@5% and sample SD for the three shared-output conditions across the ten splits.

#### Candidate prediction groups

The four-algorithm comparison used 82 shared candidate input identifiers: 28 chemical fingerprint, eight curated annotation, 22 LLM-derived annotation, ten transcriptomic perturbation, ten cell-painting and four LLM embedding inputs. In the separate MLP deployment and sensitivity analyses, the 91 candidate columns represented 44 feature sets with base and contrastive representations, plus three 64-dimensional variational-autoencoder fingerprint representations. Family counts were 31 chemical fingerprint columns, eight curated annotation columns, 22 LLM-derived annotation columns, 14 transcriptomic perturbation columns, ten cell-painting columns and six LLM embedding columns.

Among the candidate inputs, six entries labeled contrastive used raw-source predictions, and 64 configurations labeled raw used fold-fitted VAEs with 256 latent dimensions. **Supplementary Data 1** records the candidate identifiers and the representation used for each configuration.

Detailed selection groups restricted redundant predictors and were distinct from the six information families. Related SUM159 transcriptomic feature sets shared one group, and related MDA-MB-231 transcriptomic feature sets shared another. Cell-painting representations with different feature counts also shared a group. A group could contribute up to two columns, with the second restricted to the same feature set's paired representation. Combined target-and-pathway representations were limited to one column and excluded separate target or pathway representations.

#### Ensemble input selection and model settings

The response dataset contained 2,690 evaluable compounds in SUM159 and MCF7 and 2,691 in MDA-MB-231 and SK-BR-3. Ensembles were evaluated on ten repeated 80:20 train–test splits of compounds (outer splits), using seeds 12–21 and five strata based on each compound's mean PGCC response. All compared methods used the same splits. Outer training compounds were shuffled into five compound-grouped inner folds. Candidate ranking and group filtering were performed across the eligible pool separately within each inner training fold. Each family subset then retained its columns from that filtered pool.

Family subsets were scored by the unweighted mean of four cell-line Pearson correlations, averaged over the five inner validation folds. Exact ties favored fewer families and then lexicographic order. After subset selection, candidate ranking and filtering were repeated on the full outer training partition. Only columns belonging to the selected families entered the final ensemble. Validation and test missingness was filled using medians calculated from the corresponding training partition, or zero when no median was available.

TabPFN^9,10^ used a regressor with a fixed random seed. CatBoost^11^ used 250 iterations, depth four and learning rate 0.04. XGBoost^12^ used 250 trees, maximum depth four, learning rate 0.04, row and column subsampling fractions of 0.9, and L2 regularization of one. LightGBM^13^ used 250 trees, 31 leaves, learning rate 0.04, configured row and column subsampling fractions of 0.9, and L2 regularization of one.

Two-layer stacking^14^ used CatBoost, XGBoost and LightGBM as first-layer regressors. Three compound-grouped folds generated first-layer OOF predictions for the neural second layer. These predictions and cell-line indicators entered a regressor with one hidden layer of eight rectified linear units. Training used 200 epochs, mean squared error and Adam, with learning rate 0.01 and weight decay 0.0001. First-layer models were then refitted on the available training partition to predict the validation or test partition.

#### Sensitivity to separation of training stages

We compared precomputed-OOF and separated-training procedures with the same selected inputs and matched model settings.^15^

Ten repeated 80:20 train–test splits of compounds used seeds 12–21, with the same splits for both training procedures. Each split contained 2,152 training and 539 test compounds from the screening library. All available cell-line measurements for each compound remained together, and analyses used the observed responses. Configurations with and without additional annotations used fixed sets of 38 and 43 predictors, respectively (**Supplementary Data 1**). The selected gene and morphology lists, existing fingerprint-autoencoder feature tables and cellular context remained fixed.

In the precomputed-OOF procedure, base predictions were generated by five-fold cross-validation across the full compound set and then partitioned for ensemble fitting and evaluation. In separated training, five-fold OOF predictions were generated only within the ensemble-training compounds. Each base model was then fitted on all ensemble-training compounds to predict the ensemble-test compounds. Newly fitted VAE and contrastive representations and feature and response scaling used only the corresponding base-model fitting compounds. Ensemble missing-value imputation used training data only. TabPFN V3 used eight configured estimators with automatic estimator scaling and matched seeds.

Base MLPs used hidden widths of 256, 128 and 64, dropout 0.2, learning rate 0.001, weight decay 0.0001, batch size 512 and a fixed training duration of 200 epochs. These common training settings differed from the tuned base-model settings used for **Figure 5**. VAE inputs omitted the additional latent-feature scaling before MLP fitting in both procedures. A sensitivity analysis removed the ECFP6-VAE64 and PubChem-VAE64 predictors from both procedures and both ensembles across all splits; other VAE predictors and all compounds remained included.

Pearson correlation compared predicted and measured log2 PGCC count ratios within each cell line. Hits had measured count ratios ≤0.3. EF@5% divided the hit proportion among the lowest-predicted 5% by the hit prevalence in the same test set. The selected count was rounded upward to 27 compounds per cell line, with ties resolved by compound identifier. Overall means weighted cell lines equally across ten splits.

#### Performance aggregation across input families

For each family and base algorithm, feature sets were ranked by the median of their four cell-line median test-fold correlations, pooling their available representations. The two highest-ranked feature sets were selected for the family overview. For each selected family–algorithm–cell-line combination, the median and interquartile range summarize the pooled fold correlations from these feature sets and representations. For each cell line, the pooled summary contained 400 fold scores from the two selected feature sets and their representations. To rank the algorithms across all six families, we averaged the four cell-line medians within each family and then averaged the six family means. Each cell line and each family received equal weight.

#### Selection with and without expanded annotations

Selection without expanded annotations excluded the predefined GPT-derived binary target/pathway features and the subsets containing only newly added target or pathway labels. Curated annotations and the other LLM-derived features remained eligible. Seeds 12–21 defined the same ten repeated train–test splits used for the reference ensemble with expanded annotations. Each split contained 538 held-out compounds and 2,150–2,152 labeled compound–cell-line rows. Selection was performed inside each outer training partition, and comparisons with and without expanded annotations used the saved cluster predictions. Equal-cell summaries averaged the four cell-line metrics within each split before calculating the mean and sample standard deviation across splits. Pooled metrics were calculated separately across all labeled test rows.

#### External-library prediction

The external screening set comprised 21,367 LINCS and PRISM entries with at least one selected base prediction and no name or standardized parent-structure match to the PGCC screen. Additional LLM-derived target and pathway inputs were used when available. Missing selected base predictions were imputed from the corresponding training OOF medians. Input availability and configuration details are provided in **Supplementary Data 1**.

Overlap with the original PGCC screen was checked by normalized compound names or standardized parent structures against both the screened-compound catalog and the screened structural-reference table. Parent matching removed salts, neutralized charge, canonicalized tautomers and ignored stereochemistry; matches to either source were excluded. The retained point predictions were ranked in ascending order within each cell line; ties received the minimum rank. We recorded the number of available inputs for each compound.

#### Feature evaluation and gene-set analysis

Reactome gene-set enrichment used one-sided hypergeometric tests against 3,400 uniquely mapped measured CIGS genes. The LLM-prioritized genes query used 480 identifiable genes from its 481-column input. Benjamini–Hochberg correction covered all 6,684 comparisons of six lists with 1,114 eligible pathways. Pathways were selected using a fixed rule balancing coverage across lists and limiting redundant memberships; all tested pathways and exact statistics are provided in **Supplementary Data 1**.

Cell Painting feature-selection comparisons used a shared raw-feature MLP with hidden layers of 256, 128 and 64 units, dropout 0.1, learning rate 0.0005, weight decay 0.0001 and batch size 256. Training was capped at 500 epochs with early-stopping patience 80. Models included cell-line context variables and predicted PGCC and non-PGCC responses. Ten repetitions of ten-fold compound-grouped cross-validation used matched splits across configurations. Ten percent of each outer-training compound set was reserved for early stopping. Scaling parameters were estimated from the training compounds. All folds used the same fixed feature lists.

For **Fig. 3e**, the MDA-MB-231 single-cell list (29 genes) and the LLM-prioritized list (481 genes) were each compared with five fixed, response-blind random subsets of the same size from the 3,407-column CIGS matrix. All conditions used a contrastive encoder followed by the same MLP architecture and training settings. Within each outer fold, the encoder and its feature preprocessing were fitted on the fitting compounds; early-stopping and test compounds were transformed with that fitted encoder. Cellular-context features were appended after encoding. The encoder used a 512-unit hidden layer, a 128-dimensional representation and a 64-dimensional projection head, with dropout 0.1. Contrastive training used 60 epochs, batch size 256, learning rate 0.001, weight decay 0.0001 and temperature 0.1; feature dropout 0.05 and Gaussian noise with SD 0.01 generated paired views. Inputs covered 2,258 compounds, with 2,257 measured PGCC responses in SUM159 and MCF7. Ten repetitions of compound-grouped tenfold cross-validation used matched partitions across conditions. Ten percent of each outer-training compound set was reserved for early stopping. Metrics were calculated from pooled out-of-fold predictions within each cell line and repetition. A hit had a measured PGCC count ratio ≤0.30; the lowest-predicted 5% were selected, rounding upward and allocating cutoff ties fractionally. The five-size Cell Painting comparison is summarized in **Supplementary Data 1**.

Each compound was represented by six GPT-4o ratings on a 0–1 scale. Ward clustering used Euclidean distances across these six values. The dendrogram was cut into three branches. We summarized each process by its median rating within each branch. The high-rating group was the branch with the highest mean over all its compounds and all six ratings. Curated target annotations provided a descriptive check of the branches. The reported annotation summaries used exact assignments to microtubule/tubulin and Aurora kinase.

#### Precision–recall and response-error analysis

The matched evaluations of the ensemble and the training-selected individual predictor contain 21,518 test-pair appearances per method across ten splits, corresponding to 2,385 distinct compounds. Each cell-line test set has 537 or 538 compounds. For precision–recall analysis, tied prediction scores enter together; average precision is the sum of precision weighted by recall increments. AP-consistent step curves are averaged on a shared recall grid. Rank groups contain 53 or 54 compounds; exact ties are ordered by compound identity. Rank-group summaries were calculated by averaging the within-line and within-split medians equally across cell lines and then across splits.

Response-value accuracy is the fraction of predictions with absolute log2 error no greater than the specified tolerance. One log2 unit corresponds to a twofold difference in count ratio. All predictions contribute to the denominator at every tolerance. Values use equal-cell averaging within each split, followed by the mean and sample SD across ten overlapping splits.

#### External-candidate structural similarity

The structural analysis standardized molecules before computing 2,048-bit Morgan fingerprints with radius 3. Maximum Tanimoto similarity was calculated against 2,689 screened compounds with usable structures, retaining repeated standardized structures. For **Fig. 6a**, we used a nearest-neighbor Tanimoto cutoff of 0.70.

#### Supplementary Experimental Methods

PGCC and non-PGCC counts were quantified with the original screening image-analysis pipeline in SUM159, MDA-MB-231, SK-BR-3 and MCF7 cells.

Cells were allowed to attach for 24 h after seeding before treatment. Each compound and treatment condition comprised three technical wells measured on one experimental day. Voxtalisib was tested at 0.5 and 1 µM because its autofluorescence could interfere with imaging; the other compounds were tested at 1 and 10 µM. Paclitaxel (1 µM) and the test compound were added simultaneously in the combination groups. After 48 h of continuous exposure in the original treatment medium, brightfield and fluorescence images were acquired for PGCC and non-PGCC quantification.

For each cell line and plate, the mean of 12 vehicle-control wells or 12 paclitaxel-only wells provided the condition-specific denominator. PGCC and non-PGCC counts were normalized separately. The response for each compound and condition was the median of its three normalized technical wells. A median PGCC count ratio ≤0.30 was considered a hit.

For each combination-treatment well, the PGCC and non-PGCC counts were each divided by the mean count for the same population in the 12 paclitaxel-only control wells on the same plate and multiplied by 100. For each well, the PGCC reduction minus the non-PGCC reduction was calculated as the normalized non-PGCC count minus the normalized PGCC count, in percentage points. The population-specific medians and the median of the three within-well differences were calculated separately for each drug and cell line. For **Fig. 6d**, two-sided Welch's t-tests compared PGCC counts in three drug-plus-paclitaxel wells with those in 12 paclitaxel-only control wells on the same plate. For **Fig. 6e**, each well contributed log₂(PGCC count/non-PGCC count), and two-sided Welch's t-tests compared these values between the treatment and control groups. Exponentiating the mean log-ratio difference gives the treatment-to-control ratio of geometric mean PGCC/non-PGCC ratios. Confidence intervals used Welch–Satterthwaite degrees of freedom and were transformed back to the ratio scale for selectivity. Compounds on the same plate shared matched control wells. **The Validation wells sheet in Supplementary Data 1 gives raw counts for both doses and both sets of matched controls. The Figure 6 sheet gives effect estimates, 95% confidence intervals and P values for the 24 comparisons per endpoint at the reported doses.**

For the induction comparison, each plate was summarized by its mean PGCC count per well and by 100 × summed PGCC counts / summed total counts. We calculated the median of each summary across three plates.

We used the saved external-library prediction for each hit and cell line. The model was trained on compound-alone responses. For comparison with the combination experiment, both conditions used the same fixed prediction.

Within each of the four lines, Spearman correlation compared the rank order of the six hits by prediction with their order by measured median PGCC count ratio. Compound-alone measurements used vehicle controls and combination measurements used paclitaxel-only controls. One observed value per compound and cell line was obtained by summarizing the three technical wells. Zero measured counts were retained. **Supplementary Table 3** gives the descriptive rank correlations.

#### Paired bootstrap for feature-selection comparisons

For the gene-set comparison (**Fig. 3e**), each selected list was compared with the mean performance of its five fixed size-matched random lists. Pearson correlation and EF@5% were calculated separately for each model, cell line and repetition. Random-list metrics were averaged across the five lists within each repetition, and then across ten repetitions. Selected-list metrics were averaged across the same ten repetitions. The SEM was calculated as the sample SD of the ten repetition-level values divided by √10.

We evaluated the top 50, 100, 200, 300 and 500 features from the same ten-reference ranking under matched training conditions in four cell lines. The evaluation excluded the ten reference compounds, colchicine and barasertib, leaving 1,586 compounds per cell line. Within each repetition, fold-specific out-of-fold predictions were pooled to calculate Pearson correlation and top-5% enrichment in each line. Four line-specific values were averaged with equal weights. To compare 200 features with 50, 100, 300 and 500 features in four paired contrasts, we resampled the 1,586 compounds with replacement 9,999 times, applying each compound’s multiplicity to its fixed out-of-fold predictions across all five feature counts, four cell lines and ten repeats. In each draw we recalculated Pearson r and EF@5% (80 screening slots) within each line and repeat, then averaged the paired differences. Differences are reported as 200 features minus the comparator. **Supplementary Data 1** gives pointwise 95% basic bootstrap intervals and two-sided, null-centered P values for the four comparisons per endpoint.

For the gene-set comparisons, 50,000 paired compound-bootstrap draws sampled 2,258 compound identifiers with replacement, holding models and gene lists fixed. Each draw retained the same compound multiplicities across selected and random-list predictors, all available cell-line records and all ten repetitions. Missing cell-line measurements remained absent. Pearson correlation and EF@5% were recalculated within each cell line and repetition, including the screening budget, hit prevalence and cutoff ties, before averaging across repetitions. The selected-minus-random difference used the mean of the five separately calculated random-list metrics. Pointwise 95% basic intervals were twice the observed difference minus the 97.5th and 2.5th percentiles of the bootstrap differences. With B draws, the two-sided null-centered P-value was [1 + count(|Δ_boot − Δ_obs| ≥ |Δ_obs|)]/(B + 1). **Supplementary Data 1** reports the sixteen comparisons covering two lists, four cell lines and two endpoints.

#### Drug representation and algorithm comparisons

**Fig. 4g illustrates three encoder types. The ECFP6 and curated annotation examples had the largest observed mean gains within their respective sets of eight MLP-AE and 18 contrastive comparisons. The Cell Painting example comes from a separate VAE width experiment. The Figure 4g comparisons sheet in Supplementary Data 1 lists all 28 contrasts, including the fingerprint VAE comparison. Reported P values are pointwise.** For the ECFP6 fingerprint (MLP-AE) and curated targets and pathways (contrastive encoder), we averaged the four cell-line differences in held-out Pearson r within each of 100 paired folds. Two-sided paired t-tests and SEMs accounted for fold overlap across ten repeats of tenfold compound cross-validation (99 degrees of freedom).16,17 For Cell Painting, both MLPs used 200 morphology features and context variables, with or without a 64-dimensional VAE. Across ten matched holdouts, 50,000 paired compound-bootstrap draws used the same compound multiplicities across models, lines and holdouts. The SD of the mean paired changes gave the SEM, and a two-sided null-centered bootstrap gave the P value.

For the algorithm comparison in **Fig. 4i**, MLP, HGB, RF and SVM shared an eligible pool of 82 candidate inputs. Each algorithm selected its ensemble inputs on the outer training data and used the same TabPFN combiner. We compared MLP with each alternative on held-out Pearson r and EF at 5%, averaging the four cell-line values equally within each of the ten paired splits. Pointwise intervals used the two-sided overlap-corrected t statistic (nine degrees of freedom). Figure 4i stars denote the corresponding unadjusted P values for TabPFN ensembles. For **Tables 1–2**, the same test compared TabPFN with simple averaging within each base algorithm. **Supplementary Data 1** reports both ensemble endpoints.

Encoder-width sensitivity was evaluated after the screening settings were fixed. Each width was assessed by complete-ensemble hit recovery among the top-ranked 5% of compounds, with 27 selections per cell line and split across four cell lines and ten splits (1,080 testing slots). Widths used matched compound partitions within each experiment. The fingerprint experiment varied the ECFP6, MACCS and PubChem MLP-AE widths together at 16, 32, 64 and 128 dimensions. Ten affected predictors, comprising ECFP6, the three fingerprint pairings and the three-way combination in MLP and contrastive-MLP configurations, were updated. Five compound-grouped inner folds supplied their OOF training predictions; held-out predictions used the matching encoders fitted on the outer training compounds. Reconstruction fitting retained the ordered 80/20 training–validation rule and the fixed 64-unit hidden layer. The other 33 predictor columns were held fixed. The Cell Painting contrastive experiment used the same ten outer splits and 43 selected inputs, varying its encoder output at 64, 128 and 256 dimensions while holding the other 42 predictors fixed. Its encoder and MLP were fitted within five compound-grouped inner folds and then refitted on the outer training set. The encoder received 200 morphology features; context variables were appended after encoding, and the projection head remained at 64 dimensions. In both experiments, the fixed predictor columns were stored OOF predictions computed before the outer ensemble splits. TabPFN was refitted at each width using training-median imputation. The VAE experiment varied one of six screening inputs at a time across 64, 128, 256 and 512 dimensions, retaining the other predictor columns and refitting TabPFN. We averaged hit counts across the four cell lines within each split and then averaged the ten split means. The SEM was calculated as the sample SD of the ten split means divided by √10. Per-width totals, means, SEMs and split-level values are provided in **Supplementary Data 1**.

#### Cellular-context contrasts

For the individual-predictor context comparisons, **w**e sampled 2,691 compound identifiers with replacement in 20,000 draws (NumPy PCG64 seed 20261105). Each identifier's multiplicity was applied jointly to its available test records across cell lines, splits, input families and both arms. Models, input selections, predictions and split assignments were fixed. Within each split and cell line, compounds were ranked by predicted log2 PGCC count ratio, with compound identifier breaking ties; the top ceil(0.05 × n) sampled entries were selected. Duplicate sampled compounds were counted according to their multiplicity. We recalculated hit prevalence, EF@5% and Pearson correlation, averaging paired differences equally across four cell lines and ten splits. Pointwise 95% basic bootstrap intervals used twice the observed contrast minus the 97.5th and 2.5th percentiles of bootstrap contrasts. Approximate two-sided P values used the proportion of absolute null-centered bootstrap contrasts at least as large as the absolute observed contrast, with a plus-one Monte Carlo correction. Bootstrap draws with undefined EF@5% were excluded from the corresponding EF@5% summaries. Valid and excluded draw counts are reported in **Supplementary Data 1**. Relative EF@5% changes divide the paired mean difference by the mean without context. The same procedure was applied to separated versus precomputed predictions for the two primary all-selected TabPFN ensembles (EF@5% and Pearson correlation; two contrasts per endpoint). These checks used 20,000 draws with seed 202611051. **Supplementary Data 1** reports the effect estimates, P-values and pointwise intervals. Context sensitivity analysis used paired t-tests across ten matched splits with an approximate correction for overlapping training sets (**Supplementary Data 1**).

#### Correction for overlapping resamples

For each fingerprint contrast and the split-level context sensitivity analysis, let d̄ and s be the mean and sample standard deviation of the m paired differences, and q the mean of n_test/n_train across those matched resamples, counting compounds in the analyzed cohort. We calculated SE = s√(1/m + q), t = d̄/SE, and the pointwise 95% confidence interval as d̄ ± t_(0.975,m−1) × SE. We obtained two-sided P values from the t distribution with m−1 reference degrees of freedom. The mean test-to-training compound ratio was approximately 0.1111 for the fingerprint comparisons and ranged from 0.2467 to 0.2526 for the context inputs, which used ten split-level differences and nine reference degrees of freedom. The correction approximates the covariance due to overlapping resamples and uses a t approximation for the standardized mean difference.

#### Statistical comparisons for Figure 5

The evaluation used ten matched compound-level 80:20 training–test splits (seeds 12–21). The original test partitions contained 537–538 compounds per cell line. For the bootstrap, 2,385 compounds with a saved held-out prediction in at least one split formed the resampling pool. Each draw sampled 2,385 compound IDs uniformly with replacement. A single multiplicity was used for all held-out occurrences of that compound, all available cell-line records and all methods. Thus the same compounds contributed jointly across overlapping splits and paired methods. Split assignments, predictions, training-derived imputation, and model and representative selections were held fixed. Base predictions preceded ensemble splitting, and the matched retraining comparison is reported separately in **Supplementary Table 1**.

A hit had a measured PGCC count ratio ≤0.30. In each split and cell line, compounds were ordered by increasing predicted log2 count ratio. At a 5% budget, the first ceil(0.05 × n) records were selected, where n was the resampled cell-line test-set size. The original stable order resolved score ties for budget-based hit recovery. Expected random recovery was ceil(0.05 × n)/n (5.019% in the original splits). **Fig. 5g** compares each saved family-removal and reselection configuration with the complete ensemble. For **Fig. 5h**, each method separately nominated 27 compounds from the same 537–538 held-out compounds with measured responses in each cell line and split. All six families contributed one predictor in each split. Within each family, we considered its members among all 91 candidates and chose the predictor with the highest four-line mean training Pearson correlation, with alphabetical tie breaking. We used training medians for missing predictions. For each method, we averaged the number of measured hits across four cell lines within each split and then averaged the ten split means. The SEM was calculated as the sample SD of the ten split means divided by √10. Expected random hit counts were 27 times the measured hit prevalence in each test set; their SEM uses the same split-level averaging. Mean counts, SEMs and the underlying test-list values are tabulated in **Supplementary Data 1**.

For **Fig. 5c**, we compared the ensemble and individual predictor for the top 10% of ranked compounds. In each of 50,000 draws (seed 20261109), we resampled 2,385 compound IDs with replacement, preserving each ID’s multiplicity across methods, cell lines and test splits. Each method’s ranked list was rebuilt within each split and line, breaking exact score ties by compound ID. The top decile contained ceil(n/10) resampled records. We subtracted the individual-predictor median measured log2 response from the ensemble median within each cell line and split and averaged the 40 paired differences with equal weights. For decile 1, we report a pointwise basic bootstrap interval and a two-sided null-centered Monte Carlo P value. For decile 10, we report the observed difference using the same within-line and within-split medians and equal weights.

**Fig. 5e** uses complete four-line profiles (2,384 distinct held-out compounds) and the six possible line pairs per compound. Pairs were eligible when the measured responses differed by at least one log2 unit. The proportion of correctly ordered pairs was calculated within each split and then averaged across splits. A predicted tie received half credit. The cell-line mean baseline was fitted on outer-training responses.

For the comparisons in **Fig. 5a, e and g**, 50,000 paired compound-bootstrap draws (seed 20261104) produced pointwise 95% basic confidence intervals. Two-sided Monte Carlo P values were calculated as (1 + the number of null-centered differences at least as large in absolute value as the observed difference)/50,001. For **Fig. 5a**, we report the two-sided P value for the ensemble versus the training-selected individual predictor at the 5% budget. Recovery at 1% and 10% is summarized descriptively. The two comparisons of cell-line ranking accuracy in **Fig. 5e** use the same two-sided paired compound-bootstrap P values. For family-removal comparisons with non-identical predictions, we used a paired compound bootstrap and an additional two-sided paired Wilcoxon signed-rank test. **Supplementary Data 1** gives results for all five tested removals. For the Wilcoxon test, we computed the four-line mean EF@5% difference within each of ten overlapping splits, excluded zero differences, averaged ranks for tied absolute differences and enumerated all sign assignments.

#### Supplementary feature tables

**Supplementary Data 1** provides feature lists, data resources, model settings and source data for **Figures 3–6**, including both treatment conditions for all six validation compounds. **Tables 1–4** below provide the supporting comparisons and experimental measurements.

### Supplementary tables

**Supplementary Table 1**. Ensemble performance with precomputed predictions and separated training.

A. Overall means across four cell lines

| **Analysis** | **Ensemble (total inputs)** | **Pearson r Precomputed** | **Pearson r Separated** | **EF@5% Precomputed** | **EF@5% Separated** |
| --- | --- | --- | --- | --- | --- |
| All selected | With (38) | 0.302 | 0.306 | 4.23 | 4.08 |
| All selected | Without (43) | 0.298 | 0.299 | 4.38 | 4.00 |
| Two VAE inputs excluded | With (36) | 0.301 | 0.304 | 4.15 | 4.02 |
| Two VAE inputs excluded | Without (41) | 0.300 | 0.299 | 4.42 | 3.97 |

B. Cell-line means with all selected inputs

| **Expanded annotations** | **Cell line** | **Pearson r Precomputed** | **Pearson r Separated** | **EF@5% Precomputed** | **EF@5% Separated** |
| --- | --- | --- | --- | --- | --- |
| With | SUM159 | 0.302 | 0.311 | 4.18 | 3.40 |
| With | MDA-MB-231 | 0.308 | 0.310 | 2.49 | 2.41 |
| With | MCF7 | 0.254 | 0.263 | 4.66 | 5.10 |
| With | SK-BR-3 | 0.343 | 0.340 | 5.57 | 5.41 |
| Without | SUM159 | 0.297 | 0.299 | 4.26 | 3.20 |
| Without | MDA-MB-231 | 0.297 | 0.297 | 2.62 | 2.28 |
| Without | MCF7 | 0.250 | 0.258 | 4.79 | 5.00 |
| Without | SK-BR-3 | 0.350 | 0.340 | 5.87 | 5.51 |

Values are arithmetic means of split-specific metrics: ten splits and four equally weighted cell lines in A, and ten splits within each cell line in B. Precomputed denotes the global five-fold OOF procedure; separated denotes base fitting confined to ensemble-training compounds. Both use matched test compounds and settings. A hit is a measured PGCC count ratio ≤0.3; the 5% budget selects 27 compounds per cell line. The exclusion analysis removes ECFP6-VAE64 and PubChem-VAE64 from both procedures and retains all other predictors. For the primary all-selected ensembles, paired compound bootstrap comparisons gave P = 0.520 and 0.111 for EF@5% with and without expanded annotations, respectively; the corresponding P-values for Pearson correlation were 0.535 and 0.981. The bootstrap jointly resamples compound identities across cell lines and overlapping splits, conditional on saved fitted models. Exact P values and pointwise 95% intervals are reported in **Supplementary Data 1**.

**Supplementary Table 2**. Measured PGCC count ratios for the six combination-treatment hits.

| **Compound** | **Alone SUM159** | **Alone 231** | **Alone SK-BR-3** | **Alone MCF7** | **Paclitaxel SUM159** | **Paclitaxel 231** | **Paclitaxel SK-BR-3** | **Paclitaxel MCF7** |
| --- | --- | --- | --- | --- | --- | --- | --- | --- |
| AZD-8055 | **0.221** | **0.233** | **0.113** | 0.326 | **0.196** | **0.227** | **0.126** | **0.296** |
| Pyrazoloacridine | 0.498 | **0.117** | **0.036** | **0.032** | 0.367 | **0.038** | **0.103** | **0.031** |
| ONX-0914 | 0.747 | 1.389 | **0.225** | 0.780 | **0.137** | **0.169** | **0.156** | 0.464 |
| Voxtalisib† | **0.057** | **0.000** | **0.038** | **0.060** | **0.112** | **0.235** | **0.073** | **0.117** |
| Elimusertib | 1.075 | 1.625 | **0.133** | **0.289** | **0.292** | **0.252** | **0.101** | 0.305 |
| Deslanoside | 2.887 | 3.000 | **0.209** | 0.590 | **0.285** | **0.218** | **0.096** | 0.503 |

Values are medians of three technical wells, normalized to the same-plate vehicle control (alone) or paclitaxel-only control. Bold values meet the hit cutoff ≤0.30; classification uses full precision. 231 denotes MDA-MB-231. †Voxtalisib was at 0.5 µM; other compounds were at 10 µM and paclitaxel at 1 µM. In paclitaxel-treated cultures, PGCC counts were lower than matched paclitaxel-only controls in all 24 compound–cell-line comparisons (two-sided Welch's t-tests, all P < 0.001; three treatment wells and 12 control wells per comparison). The within-well log₂(PGCC/non-PGCC) ratio was lower than control in 23 of 24 comparisons at P < 0.05; ONX-0914 in MDA-MB-231 had P = 0.1068. **Supplementary Data 1** provides the individual treatment and control wells, effect estimates, 95% confidence intervals and exact P-values.

**Supplementary Table 3**. Within-line agreement between predicted response and measured response for the six combination-treatment hits.

| **Cell line** | **Compounds** | **Technical wells per value** | **ρ, drug alone** | **ρ, with PTX** |
| --- | --- | --- | --- | --- |
| SUM159 | 6 | 3 | 0.71 | 0.49 |
| MDA-MB-231 | 6 | 3 | 0.77 | 0.14 |
| SK-BR-3 | 6 | 3 | -0.54 | -0.31 |
| MCF7 | 6 | 3 | -0.66 | -0.54 |

Spearman ρ compares the same six saved compound-alone predictions with each measured condition within a cell line. Values are rank correlations among compounds selected by the measured combination-treatment hit criterion. PTX denotes paclitaxel.

**Supplementary Table 4**. Enrichment with drug inputs alone, cell-line identity or molecular context.

| **Drug-feature family** | **Drug only** | **Drug + cell-line ID** | **Drug + molecular context** |
| --- | --- | --- | --- |
| Chemical fingerprints | 2.35 ± 0.74 | 1.90 ± 0.35 | 2.27 ± 0.45 |
| Curated annotations | 2.36 ± 0.48 | 2.87 ± 0.32 | 2.75 ± 0.31 |
| LLM-derived annotations | 2.97 ± 0.39 | 3.10 ± 0.40 | 3.11 ± 0.37 |
| Text embeddings | 2.94 ± 0.86 | 2.34 ± 0.41 | 2.84 ± 0.44 |
| Cell Painting | 2.18 ± 1.23 | 1.94 ± 0.37 | 2.19 ± 0.94 |
| Transcriptomic perturbation | 1.88 ± 0.71 | 2.17 ± 0.44 | 2.18 ± 0.46 |

Values are EF@5% means ± sample SD across ten matched 80:20 compound splits, using one fixed drug input per family. All three conditions use a shared output layer predicting PGCC and non-PGCC responses. Within each split, enrichment is calculated separately for SUM159, MDA-MB-231, MCF7 and SK-BR-3 and averaged equally across the four lines. Model settings are described under Cell-line identity comparison.

#### Supplementary Figure 1 Effects of cellular context on individual predictors

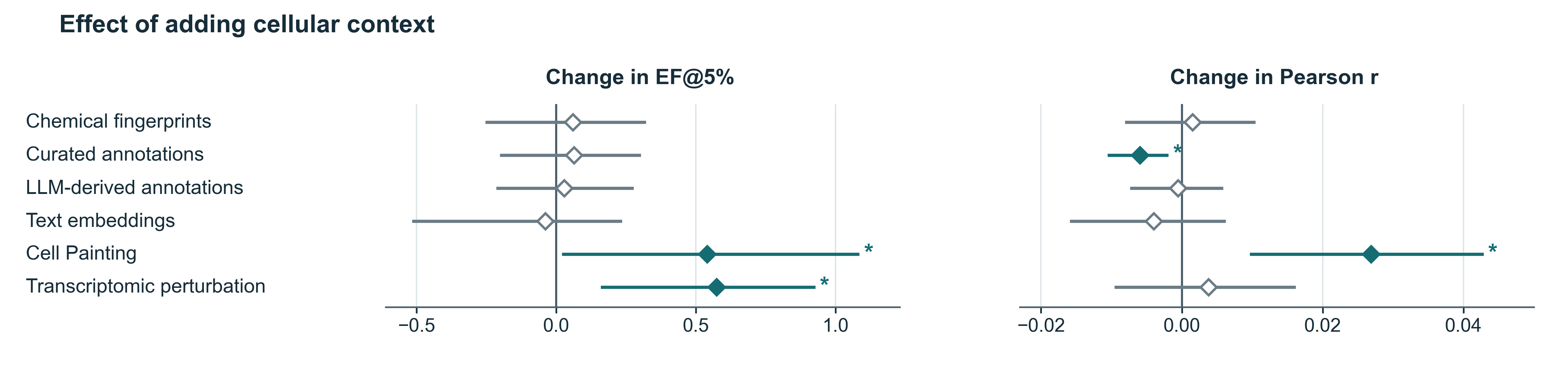

**Supplementary Figure 1. Effects of cellular context on individual predictors.** Paired MLP predictors used the same drug inputs and cell-line-specific output layers, with or without molecular cellular-context features. Points show mean differences in EF@5% (left) and Pearson correlation (right), calculated as with minus without context. Values are averaged equally across four cell lines and ten matched compound splits. Whiskers are pointwise 95% basic intervals from 20,000 paired compound-bootstrap draws. Filled diamonds and asterisks indicate two-sided compound-bootstrap P < 0.05. The no-context condition set standardized cellular-context coordinates to zero during fitting and prediction. Full settings and resampling definitions are given in **Supplementary Computational Methods**; exact results are in **Supplementary Data 1**.

#### Supplementary Figure 2 Sensitivity to VAE width across six screening inputs

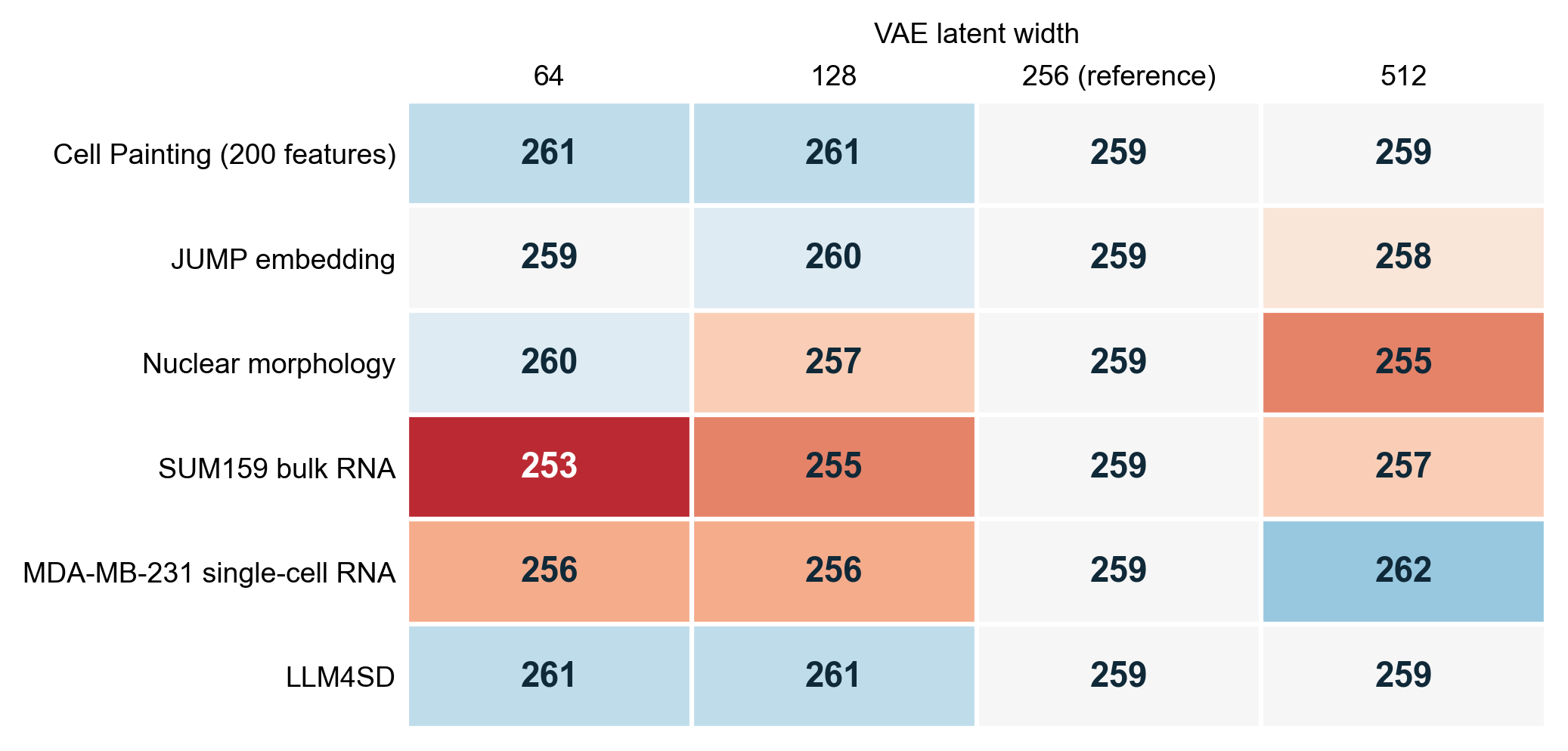

**Supplementary Figure 2**. Measured hits recovered by the ensemble when changing the VAE width of one input. Each row replaces one predictor’s 256-dimensional representation with 64, 128 or 512 dimensions and holds the other inputs fixed. The evaluated deployment configuration excluded additional LLM-derived binary target and pathway labels. Each width used the same ten compound splits and four cell lines, with 27 selections per cell line and split (1,080 screening slots). An appearance counted as a hit when the measured PGCC count ratio was ≤0.30. A compound could contribute in multiple lists. Blue and red indicate more and fewer recovered hits, respectively, than the 256-dimensional reference.

### Supplementary references

1. Chen, T., Kornblith, S., Norouzi, M. & Hinton, G. A simple framework for contrastive learning of visual representations. Proc. Mach. Learn. Res. **119**, 1597–1607 (2020). https://proceedings.mlr.press/v119/chen20j.html.

2. Kingma, D. P. & Welling, M. Auto-encoding variational Bayes. Preprint at https://doi.org/10.48550/arxiv.1312.6114 (2013).

3. Chandrasekaran, S. N. et al. Three million images and morphological profiles of cells treated with matched chemical and genetic perturbations. *Nat. Methods* **21**, 1114–1121 (2024). https://doi.org/10.1038/s41592-024-02241-6.

4. Benjamini, Y. & Hochberg, Y. Controlling the false discovery rate: a practical and powerful approach to multiple testing. *J. R. Stat. Soc. B* **57**, 289–300 (1995). https://doi.org/10.1111/j.2517-6161.1995.tb02031.x.

5. Zdrazil, B. et al. The ChEMBL Database in 2023: a drug discovery platform spanning multiple bioactivity data types and time periods. *Nucleic Acids Res.* **52**, D1180–D1192 (2024). https://doi.org/10.1093/nar/gkad1004.

6. Cannon, M. et al. DGIdb 5.0: rebuilding the drug-gene interaction database for precision medicine and drug discovery platforms. *Nucleic Acids Res.* **52**, D1227–D1235 (2024). https://doi.org/10.1093/nar/gkad1040.

7. UniProt Consortium. UniProt: the Universal Protein Knowledgebase in 2025. *Nucleic Acids Res.* **53**, D609–D617 (2025). https://doi.org/10.1093/nar/gkae1010.

8. Milacic, M. et al. The Reactome Pathway Knowledgebase 2024. *Nucleic Acids Res.* **52**, D672–D678 (2024). https://doi.org/10.1093/nar/gkad1025.

9. Hollmann, N. et al. Accurate predictions on small data with a tabular foundation model. *Nature* **637**, 319–326 (2025). https://doi.org/10.1038/s41586-024-08328-6.

10. Grinsztajn, L. et al. TabPFN-3: technical report. Preprint at https://doi.org/10.48550/arxiv.2605.13986 (2026).

11. Prokhorenkova, L., Gusev, G., Vorobev, A., Dorogush, A. V. & Gulin, A. CatBoost: unbiased boosting with categorical features. Adv. Neural Inf. Process. Syst. **31** (2018). https://proceedings.neurips.cc/paper/2018/hash/14491b756b3a51daac41c24863285549-Abstract.html.

12. Chen, T. & Guestrin, C. XGBoost: a scalable tree boosting system. In Proc. 22nd ACM SIGKDD International Conference on Knowledge Discovery and Data Mining 785–794 (2016). https://doi.org/10.1145/2939672.2939785.

13. Ke, G. et al. LightGBM: a highly efficient gradient boosting decision tree. Adv. Neural Inf. Process. Syst. **30** (2017). https://proceedings.neurips.cc/paper/2017/hash/6449f44a102fde848669bdd9eb6b76fa-Abstract.html.

14. Wolpert, D. H. Stacked generalization. *Neural Netw.* **5**, 241–259 (1992). https://doi.org/10.1016/s0893-6080(05)80023-1.

15. Mnich, K., Kitlas Golińska, A., Polewko-Klim, A. & Rudnicki, W. R. Bootstrap bias corrected cross validation applied to super learning. In Computational Science – ICCS 2020. Lect. Notes Comput. Sci. **12139**, 550–563 (2020). https://doi.org/10.1007/978-3-030-50420-5_41.

16. Bouckaert, R. R. & Frank, E. Evaluating the replicability of significance tests for comparing learning algorithms. In Advances in Knowledge Discovery and Data Mining 3–12 (2004). https://doi.org/10.1007/978-3-540-24775-3_3.

17. Nadeau, C. & Bengio, Y. Inference for the generalization error. Mach. Learn. 52, 239–281 (2003). https://doi.org/10.1023/A:1024068626366.
